# Comprehensive quantitative proteome analysis of *Aedes aegypti* identifies proteins and pathways involved in *Wolbachia pipientis* and Zika virus interference phenomenon

**DOI:** 10.1101/2020.12.12.422506

**Authors:** Michele Martins, Luis Felipe Costa Ramos, Jimmy Rodriguez Murillo, André Torres, Stephanie Serafim de Carvalho, Gilberto Barbosa Domont, Danielle Maria Perpétua de Oliveira, Rafael Dias Mesquita, Fábio César Sousa Nogueira, Rafael Maciel-de-Freitas, Magno Junqueira

## Abstract

Zika virus is a global public health emergency due to its association with microcephaly, Guillain-Barré syndrome, neuropathy, and myelitis in children and adults. A total of 87 countries have had evidence of autochthonous mosquito-borne transmission of Zika virus, distributed across four continents, and no antivirus therapy or vaccines are available. Therefore, several strategies have been developed to target the main mosquito vector, *Aedes aegypti*, to reduce the burden of different arboviruses. Among such strategies, the use of the maternally-inherited endosymbiont *Wolbachia pipientis* has been applied successfully to reduce virus susceptibility and decrease transmission. However, the mechanisms by which *Wolbachia* orchestrate resistance to ZIKV infection remain to be elucidated. In this study, we apply isobaric labeling quantitative mass spectrometry-based proteomics to quantify proteins and identify pathways altered during ZIKV infection; *Wolbachia* infection; co-infection with *Wolbachia/*ZIKV in the *Ae. aegypti* heads and salivary glands. We show that *Wolbachia* regulates proteins involved in ROS production, regulates humoral immune response, and antioxidant production. The reduction of ZIKV polyprotein in the presence of *Wolbachia* in mosquitoes was determined by mass spectrometry and corroborates the idea that *Wolbachia* helps to block ZIKV infections in *Ae. aegypti*. The present study offers a rich resource of data that may help to elucidate mechanisms by which *Wolbachia* orchestrate resistance to ZIKV infection in *Ae. aegypti*, and represents a step further on the development of new targeted methods to detect and quantify ZIKV and *Wolbachia* directly in complex tissues.

**Highlights:**

- The abundance of ZIKV polyprotein is reduced in the presence of *Wolbachia*
- Shotgun proteomics quantifies ZIKV and *Wolbachia* proteins directly in tissues
- *Wolbachia* regulates proteins involved in ROS production
- *Wolbachia* regulates humoral immune response and antioxidant production
- Metabolism and detoxification processes were associated with mono infections

## 1 Introduction

Zika virus (ZIKV) is a single-stranded RNA virus that belongs to the *Flavivirus* genus and *Flaviviridae* family. Originally discovered in a primate from the Zika forest in Uganda in 1947, remained unimportant for public health until the beginning of the XX century (Dick et al., 1952). In 2014, ZIKV emerged in the Pacific islands and a few years later invaded the Americas, leading the World Health Organization to claim a global public health emergency due to its association with microcephaly in newborns and Guillain-Barré syndrome, neuropathy, and myelitis in children and adults (Campos et al., 2015; Barreto et al., 2016). As of May 2017, around 75% of inhabitants of the Yap Islands were infected with ZIKV, but no microcephaly was recorded. Whilst in Brazil more than 220,000 cases were recorded and more than 2,600 newborns had birth defects like microcephaly and other neuropathies (Baud et al., 2017).

The mosquito *Aedes aegypti* is the primary vector of ZIKV worldwide (Chouin-Carneiro et al., 2016; Ferreira-de-Brito et al., 2016). To date, no antivirus therapy or vaccines are available to mitigate ZIKV transmission, which increases the urgency for effective methods targeting the mosquito vector to reduce the burden of arboviruses as ZIKV. Among such strategies, the use of the maternally-inherited endosymbiont *Wolbachia pipientis* has gained attention recently. The bacterium *Wolbachia* is naturally found in around 60% of arthropod species. Although *Wolbachia* does not naturally infect *Ae. aegypti*, some strains were transinfected by microinjection into this mosquito species with the further discovery of their pathogen-blocking properties (Moreira et al., 2009; Aliota et al., 2016; Dutra et al., 2016). The rationale underlying a mass release of *Wolbachia* infected mosquitoes is to replace the local *Ae. aegypti* population. Usually, it has high vector competence to arboviruses, like ZIKV, while the *Wolbachia* transinfected mosquitoes have reduced susceptibility and decreased transmission (Hoffmann et al., 2011). This approach is currently ongoing in at least 12 countries and the first communications showing its impact on disease transmission decrease are being revealed (Nazni et al., 2019; Indriani et al., 2020; Ryan et al., 2020).

The mechanisms which are orchestrated by *Wolbachia*, conferring resistance to arboviruses infection remain to be elucidated, however, it has been assigned to the activation of the mosquito immune system, down-regulation of genes that encode proteins or receptors necessary to the virus binding and depletion of host’s resources essentials to the virus life cycle (Ford et al., 2020). When infected with *Wolbachia*, the mosquito Toll pathway is activated through the increase of reactive oxygen species (ROS), resulting in the production of antimicrobial peptides like cecropin (Pan et al., 2012). The gene expression modulation of tubulin, insulin receptors, and other genes seems to hinder the virus infection (Haqshenas et al., 2019; Paingankar et al., 2020). Also, it has been proposed that the lipids are relevant to *Wolbachia* infection. While one study implies it competes with the host’s lipid resources (mainly cholesterol), others suggest it modulates the lipid production which can have a role blocking virus replication (Geoghegan et al., 2017; Koh et al., 2020; Manokaran et al., 2020).

Protein quantification on a genome-wide scale allows a systemic bird’s-eye view of protein variation and pathway regulation under different conditions and samples. (Choudhary and Mann, 2010). Therefore, it helps to highlight some features and emergent properties of complex systems, which could be obscured when the analysis is performed only from a reductionist point of view. In this way, mass spectrometry (MS)-based proteomics, together with robust bioinformatic tools, allows the formulation of more ambitious hypotheses due to the possibility of generating large and detailed datasets. Reliable large-scale protein quantification can be achieved using isobaric labeling and shotgun proteomics (Pappireddi et al., 2019). Several quantitative methods have already been successfully applied in insects to access differentially regulated proteins (De Mandal et al., 2020; García-Robles et al., 2020; Serteyn et al., 2020). The *Ae. aegypti* infection with Mayaro virus or microsporidian parasites have already been studied (Duncan et al., 2012; Vasconcellos et al., 2020) and a previous descriptive work was based on comparing the proteome of *Ae. aegypti* female heads during blood and sugar meals conditions (Nunes et al., 2016).

Here, we use an isobaric labeling-based quantitative proteomic strategy to interrogate the interaction between the *Wolbachia w*Mel strain and ZIKV infection in *Ae. aegypti* heads and salivary glands. *Wolbachia* is present in several tissues and organs in *Ae. aegypti* mosquitoes, with higher density in ovaries and tissues on the head and anterior part of the thorax, like the ommatidia cells, brain, salivary glands and cardia (Moreira et al., 2009). To the best of our knowledge, this is the most complete proteome of *Ae. aegypti* reported so far, with a total of 3790 proteins identified, corresponding to 25.75% of total protein-coding genes. This work includes the identification and quantification of several peptides from the ZIKV and 323 *Wolbachia* proteins in a complex sample tissue, paving the way to the development of sensitive approaches to detect and quantify the ZIKV and *Wolbachia* directly in the *Ae. aegypti* tissue via mass spectrometry. Moreover, we describe and discuss proteins and pathways altered in *Ae. aegypti* during ZIKV infections, *Wolbachia* infections, co-infection *Wolbachia*/ZIKV, and compared with no infection.

## 2 Material and methods

### 2.1 Mosquitoes

To access the effects of ZIKV and *Wolbachia* infection on *Ae. aegypti* proteome, we used mosquitoes from two different sites in Rio de Janeiro (Rio de Janeiro State, Brazil): Tubiacanga (22°47’06”S; 43°13’32”W) and Porto (22°53′43″ S, 43°11′03″ W), distant 13km apart from each other. Tubiacanga was selected to represent an area in which mosquitoes have *Wolbachia* (*w*Mel strain) since it is the first site in Latin America with an established invasion (>90% frequency) (Garcia et al., 2019). Porto represents a *Wolbachia* free area. Eggs were collected through 80 ovitraps (Codeço et al., 2015) installed every 25-50 m each other in both sites. Ovitraps were placed over an extensive geographic area to ensure we captured the local *Ae. aegypti* genetic variability, collecting at least 500 eggs per site. The eggs were hatched and the mosquitoes were maintained at the insectary under a relative humidity of 80 ± 5% and a temperature of 25 ± 3°C, with *ad libitum* access to a 10% sucrose solution. The experimental infection was performed with mosquitoes from the F1 generation.

### 2.2 Viral strain

*Aedes aegypti* females were orally infected with the ZIKV strain Asian genotype isolated from a patient in Rio de Janeiro (GenBank accession number KU926309). Previous reports have shown high infectivity of this ZIKV isolate to Rio de Janeiro *Ae. aegypti* populations, including Porto, which had 100% infected females after receive an infective blood meal (Fernandes et al., 2016; da Silveira et al., 2018; Petersen et al., 2018). Viral titers in supernatants were previously determined by serial dilutions in Vero cells, expressed in plaque-forming units per milliliters (PFU/ml). All the assays were performed with samples containing 3.55 × 10^6^ PFU/ml. Viral stocks were maintained at −80°C until its use.

### 2.3 ZIKV infection

Sugar supply was removed 36-h before mosquitoes were challenged with the infective blood meal to increase female’s avidity. Six-seven days old inseminated *Ae. aegypti* females from each of the two populations (Tubiacanga and Porto) were separated in cylindrical plastic cages for blood-feeding. Oral infection procedures were performed through a membrane feeding system (Hemotek, Great Harwood, UK), adapted with a pig-gut covering, which gives access to human blood. The infective blood meals consisted of 1 ml of the supernatant of infected cell culture, 2 ml of washed rabbit erythrocytes, and 0.5 mM of adenosine triphosphate (ATP) as phagostimulant. The same procedure and membrane feeding apparatus were used to feed control mosquitoes, but they received a noninfectious blood meal, with 1 ml of cell culture medium replacing the viral supernatant. After the experimental infection, we had a total of 152 *Ae. aegypti* females.

### 2.4 Ethical approval

The maintenance of *Ae. aegypti* colonies with *Wolbachia* in the lab were achieved by offering blood obtained from anonymous donors from the blood bank of the Rio de Janeiro State University, whose blood bags would be discarded due to small volume. We have no information on donors, including sex, age, and clinical condition, but the blood bank discards those bags positive for Hepatitis B, Hepatitis C, Chagas disease, syphilis, human immunodeficiency virus, and human T-cell lymphotropic virus. Before offering the blood for mosquitoes, it was screened for Dengue virus (DENV) using the Dengue NS1 Ag STRIP (Bio-Rad). The use of human blood was approved by the Fiocruz Ethical Committee (CAAE 53419815.9.0000.5248).

### 2.5 Protein extraction for proteomics

Proteins from a total of 152 *Ae. aegypti* females were processed, of which 39 were coinfected with *Wolbachia* and ZIKV (WZ); 36 non-infected (A); 35 infected with *Wolbachia* (W) only and 42 infected with ZIKV (Z). On 14 days post-infection, each mosquito head plus the salivary gland was separated from the body using needles and a scalpel blade which were sterilized after every single use (Schmid et al., 2017). Proteins were extracted by lysis with a buffer containing 7 M urea, 2 M thiourea, 50 mM HEPES pH 8, 75 mM NaCl, 1 mM EDTA, 1 mM PMSF, and protease/phosphatase inhibitor cocktails (Roche). Then, samples were vortexed, sonicated in a cold bath for 10 minutes and sonicated by the probe for 10 minutes. Lysates were centrifuged at 10,000 *g* for 10 minutes at 4°C and the supernatants were carefully transferred to new tubes for further steps.

### 2.6 Protein digestion and iTRAQ labeling

The protein concentration was determined by the Qubit Protein Assay Kit® fluorometric (Invitrogen), following the manufacturer’s instructions. A total of 100 μg of proteins from each condition was processed. Protein samples were incubated with dithiothreitol (DTT -GE Healthcare) at 10 mM and 30°C for one hour. Subsequently, iodoacetamide solution (GE Healthcare) was added, with a final concentration of 40 mM. The reaction was performed at room temperature, in the dark, for 30 minutes. The samples were diluted 10x with 50 mM HEPES buffer to reduce the concentration of urea/thiourea. The solution was incubated with trypsin (Promega) in a 1:50 (w/w, enzyme/protein) ratio at 37° C for 18 hours. The resulting peptides were desalted with a C-18 macro spin column (Harvard Apparatus) and then vacuum dried. The dried peptides were labeled with isobaric tags for relative and absolute quantitation (iTRAQ) 4-plex (ABSciex). The labeling process was performed according to Murillo et al. (2017). Briefly, dried peptides were dissolved in 50 mM TEAB and iTRAQ tags were dissolved in dry ethanol, labeling was performed by mixing peptide and tag solutions for one hour at room temperature. The four tag reactions were combined in a single tube (ratio 1:1:1:1) and dried in a speed vac prior to fractionation. Each condition was labeled as follows i) Tag 114 corresponds to sample W (*Wolbachia* infected); ii) Tag 115 to sample A (none infection); iii) Tag 116 to sample WZ (*Wolbachia* and ZIKV co-infection); and iv) Tag 117 to sample Z (ZIKV infection).

### 2.7 Fractionation by hydrophilic interaction chromatography and LC-MS/MS analysis

The iTRAQ labeled peptide mixture was fractionated off-line by HILIC (Hydrophilic interaction liquid chromatography) before LC-MS/MS analysis. The dried samples were resuspended in acetonitrile (ACN) 90% / trifluoroacetic acid (TFA) 0.1% and injected into Shimadzu UFLC chromatography using a TSKGel Amide-80 column (15 cm x 2 mm i.d. x 3 μm - Supelco), flow rate 0.2 ml/minute; mobile phases A (ACN 85%/TFA 0. 1%) and B (TFA 0.1%); gradient: 0% of phase B in 5 minutes; 0% to 10% of phase B from 5 to 10 minutes; 10% to 20% of phase B from 10 to 55 minutes; 20% to 100% of phase B from 55 to 60 minutes; and 100% to 0% of phase B from 60 to 65 minutes. For each pooled sample, 8 fractions were collected and combined according to the separation and intensity of the peaks. The pools of fractions were dried in a speed vac and resuspend in 0.1% formic acid (FA). The samples were analyzed in Easy-nLC 1000 coupled to a Q-Exactive Plus mass spectrometer (Thermo Scientific). For each fraction (or pool of fractions), a linear gradient of 5% to 40% of B em 105 min, 40-95% in 8 min and 95% B in 5 min was performed in 120 minutes; phase A (0.1% FA), B (ACN 95%, 0.1% FA). Ionization was performed in an electrospray source with the acquisition of spectra in positive polarity by data-dependent acquisition (DDA) mode, spray voltage of 2.5 kV, and temperature of 200°C in the heated capillary. The acquisition was set as follows: full scan or MS1 in the range of 375 - 1800 m/z, resolution of 70,000 (m/z 200), fragmentation of the 10 most intense ions in the HCD collision cell, with standardized collision energy (NCE) of 30, resolution of 17,000 in the acquisition of MS/MS spectra, the first mass of 110 m/z, isolation window of 2.0 m/z and dynamic exclusion of 45 s.

### 2.8 Data analysis

The raw data were processed using Proteome Discoverer 2.4 Software (Thermo Scientific). Peptide identification was performed with the Sequest HT search engine against *Ae. aegypti* (genome version/assembly ID: INSDC: GCA_002204515.1), ZIKV, and *Wolbachia* databases provided by VectorBase (Giraldo-Calderón et al., 2015), ViPR (Pickett et al., 2012), and UniProt (The UniProt Consortium, 2019) respectively. Searches were performed with peptide mass tolerance of 10 ppm, MS/MS tolerance of 0.1 Da, tryptic cleavage specificity, two maximum missed cleavage sites allowed, fixed modification of carbamidomethyl (Cys) and variable modification of iTRAQ 4-plex (Tyr, Lys and peptide N-terminus), phosphate (Ser, Thr and Tyr) and oxidation (Met); peptides with high confidence were considered. False discovery rates (FDR) were obtained using Target Decoy PSM selecting identifications with a q-value equal or to less than 0.01. Data of all technical replicates were log2-transformed and normalized by subtracting the column median. Relative iTRAQ quantification analysis was carried out in the Perseus software, version 1.6.12.0 (Tyanova et al., 2016), based on the intensity of the fragmented reporter peaks in MS / MS.

### 2.9 Statistics and gene enrichment

For ZIKV peptides abundance comparison, statistical analysis was performed in experimental groups using T-test analysis (95% confidence values, p<0.05), meanwhile for differential proteins was used one-way analysis of variance (ANOVA) and Tukey’s multiple comparisons post-test (95% confidence values, p<0.05) in Graphpad Prism 8.0.0 for Windows, GraphPad Software, San Diego, California USA, www.graphpad.com. Biological Processes enrichment for differentially expressed proteins were performed using Fisher’s Exact Test (p<0.05) in VectorBase website (bit.ly/38OmEX0) (Giraldo-Calderón et al., 2015; Ashburner et al., 2000). The resulting gene ontology (GO) lists were summarized and represented in a network interaction format in REVIGO (Supek et al., 2011).

## 3. Results and discussion

### 3.1 Proteome analysis

Proteomics has improved in recent years, especially considering its coverage and sensitivity, which offer more opportunities to study changes in the global proteome of complex tissues (Aebersold and Mann, 2016). Therefore, the use of isobaric-labeled quantitative proteomics measurements provides efficiency and increased depth coverage of proteomics analysis (Rauniyar and Yates, 2014). Peptides from the four experimental groups were labeled with iTRAQ 4-plex, enabling multiplexing of samples prior to off-line fractionation and LC-MS/MS analysis (Fig. 1). We were able to identify 4,117 protein groups (Supplementary Table 1), 26,898 peptides and 909,046 MS/MS spectra. The protein search was performed using *Ae. aegypti*, ZIKV, and *Wolbachia* databases simultaneously. Of all identified proteins, 3,790 belong to *Ae. aegypti*, 323 to *Wolbachia* and 3 unique peptides with a total of 10 PSMs to the ZIKV polyprotein (Fig. 2, Supplementary Figure 1, and Supplementary Figure 2).

**Figure 1:**
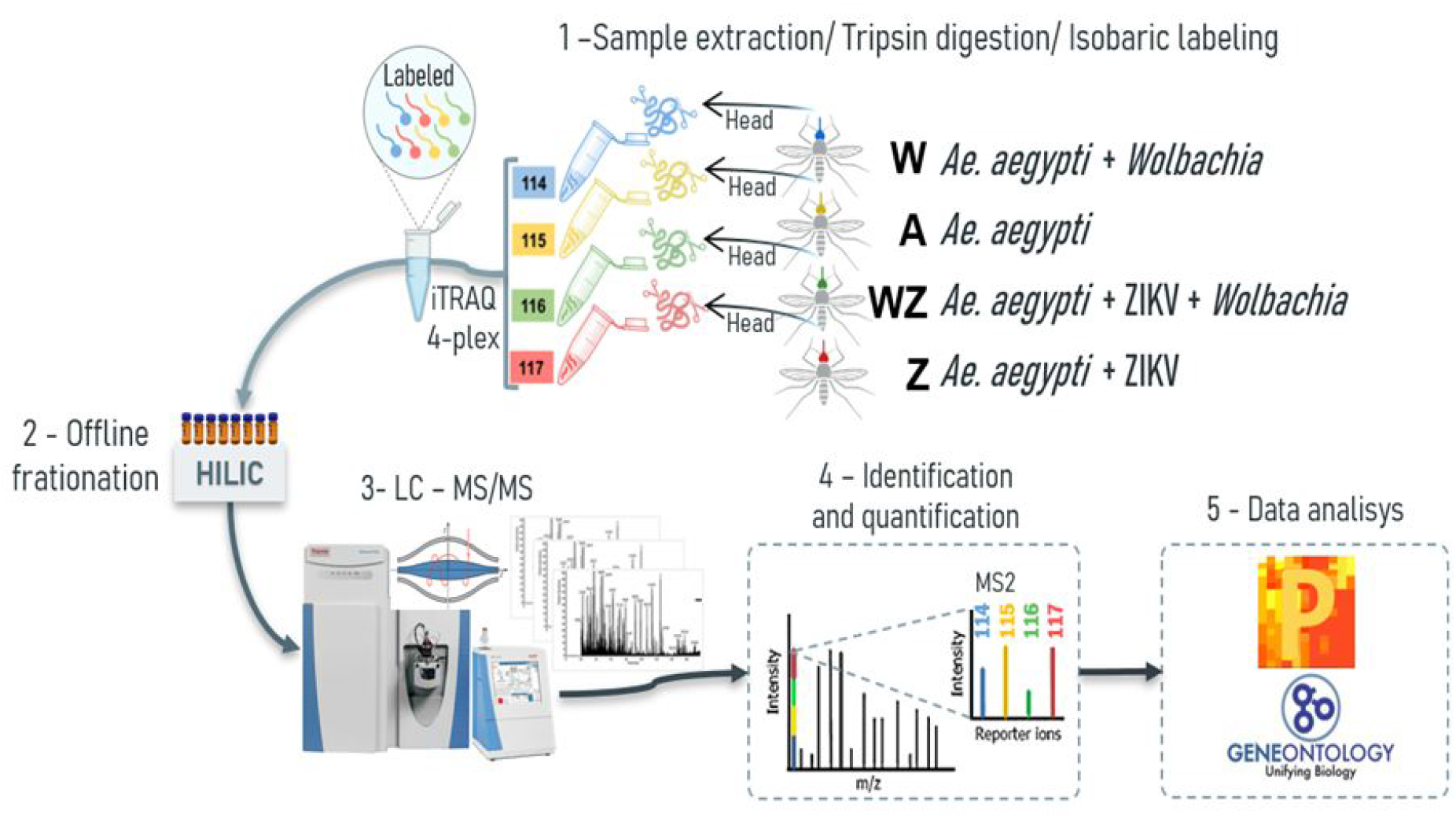
The proteomics workflow carried out in the head+salivary gland of *Ae. aegypti* females used samples from four different conditions: A - non-infected mosquitoes; W - *Wolbachia* infected; Z - ZIKV infected; and WZ - *Wolbachia* and ZIKV infection. **1** - The samples were subjected to lysis and, subsequently, proteins were reduced, alkylated, and digested using trypsin. The resulting peptides were labeled with iTRAQ-4plex and combined in 1:1:1:1 ratio; **2** - Labeled peptides were fractionated offline using HILIC chromatography; **3** - Pool of fractions were analyzed by nLC–MS/MS in Q-Exactive Plus mass spectrometer; **4** - Fragmentation of the attached tag release a low *m/z* reporter ions used for relative quantification of the peptides/proteins from which they originated; **5**-Statistical analysis were performed in Perseus and bioinformatics analysis used Gene Ontology (GO) in the enrichment of proteins differentially regulated.

**Figure 2:**
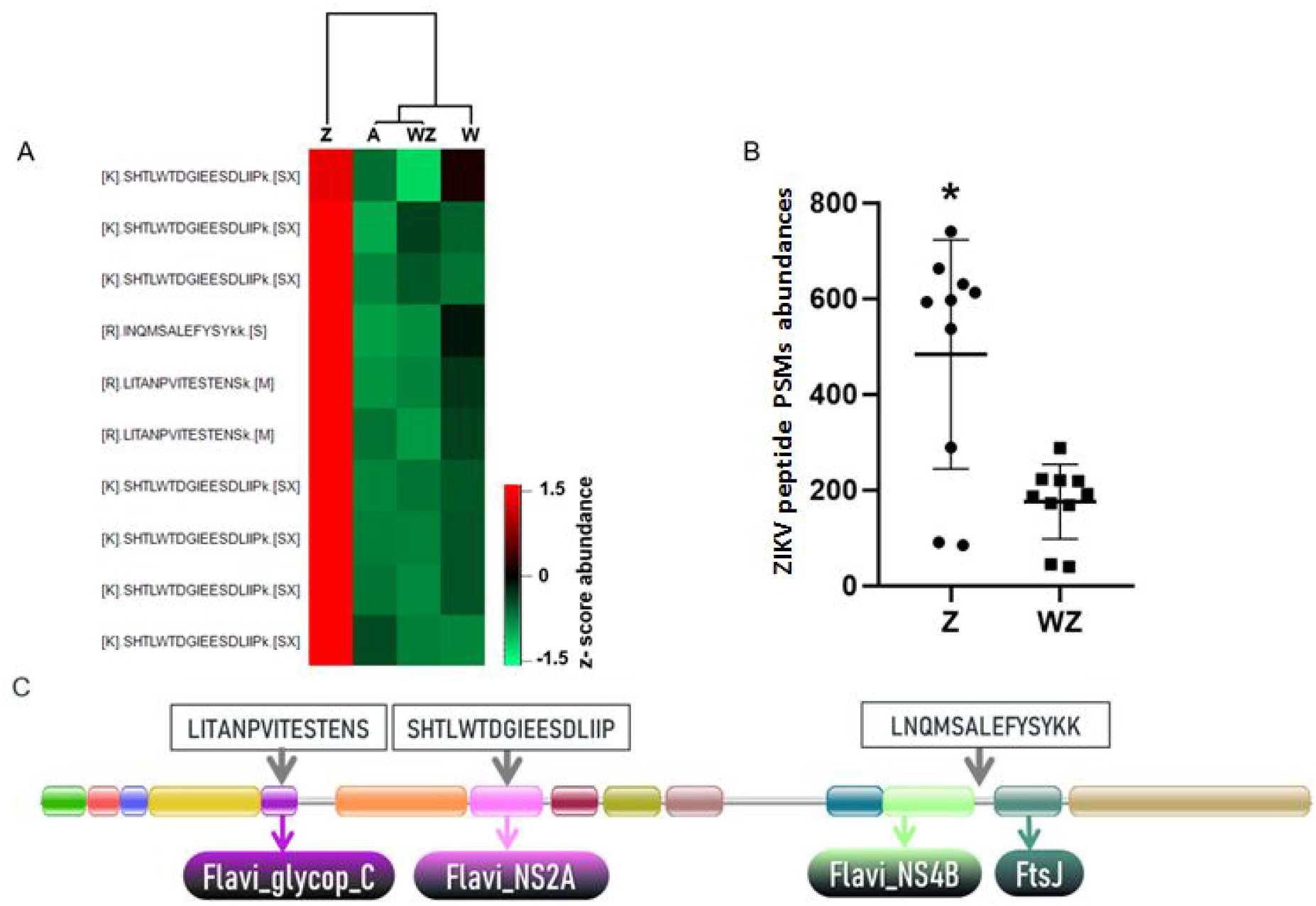
Identification and quantification of ZIKV peptides. A: Heatmap of ZIKV PSMs identified in the *Ae. aegypti* head+salivary gland generated in Perseus software using abundance values transformed by z-score. B: Total abundance of peptides spectrum matches (PSMs) calculated from report ions intensities from the three ZIKV peptides identified in samples. Statistical analysis (T-test; * p-value 0.0011) was developed by using the software GraphPad Prism 7.0. C: Domain architecture of the complete ZIKV polyprotein obtained from the PFAM database (PF01570) (El-Gebali et al., 2018). The location within the conserved domains and amino acid sequence of the three unique ZIKV peptides identified in proteomics analysis are highlighted.

### 3.2 Quantitative proteomics

The quantitative proteomics analysis was performed by comparing the four experimental groups using the normalized iTRAQ reporter ion intensity of identified peptides. Replicates were highly correlated (Pearson correlation mean of 0.97) and normalization methods were applied (Supplementary figure 3). The 196 proteins with significant differences were statistically determined by the ANOVA test. Using the Tukey post-test ANOVA, we defined pairs of proteins with significant differences between the groups (Supplementary table 2). The comparisons between the samples were: single-infected mosquitoes versus non-infected mosquitoes (W versus A and Z versus A); coinfected versus non-infected mosquitoes (WZ versus A); coinfected versus single-infected mosquitoes (WZ versus W and WZ versus Z).

#### 3.2.1 Abundance of ZIKV polyprotein is reduced in the presence of *Wolbachia*

The quantitative analysis using the iTRAQ reporter ions from ZIKV peptides revealed that ZIKV proteins were more abundant in the absence of *Wolbachia* in contrast to mosquitoes coinfected with ZIKV and *Wolbachia* (Fig 2). This latter condition showed reporter ions intensity compared to the background signal of uninfected mosquitoes (Fig 2A). This result indicates *Wolbachia* presence likely reduces virus replication in the female head and salivary gland (Fig 2B). Noteworthy, the presence of the *w*Mel strain in organs like salivary glands and ovaries of *Ae. aegypti* females induce the antiviral response that supports *Wolbachia* deployment to mitigate disease transmission.

The detection of ZIKV in the mosquito head + salivary gland was possible even considering that virus proteins were a minute fraction of all proteins identified. Peptides LITANPVITESTENS, SHTLWTDGIEESDLIIP, and LNQMSALEFYSYKK showed a robust increase in ZIKV infected tissue and this data represents a resource to further develop targeted proteomics methods to quantify ZIKV in infected mosquitoes (MS / MS spectra shown in Supplementary Figure 1).

#### 3.2.2 Identification of salivary gland proteins, neuropeptides and hormones in head samples

During a blood meal, the female mosquitoes inject saliva into the host skin. The salivary glands produce a huge amount of molecules to avoid host responses against blood loss and to evade the host immune system (Ribeiro et al., 2016). For transmission of arboviroses, the virus must be secreted with the saliva during a blood meal in a vertebrate host (Linthicum et al., 1996). In this work, we took advantage of the sensitivity of protein detection by MS to detect and quantify proteins from the salivary gland, which is located in the heads of *Ae. aegypti*. We compared the head proteome with two proteomes and one transcriptome (Wasinpiyamongkol et al., 2010; Dhawan et al., 2017; Ribeiro et al., 2016) from salivary glands of female mosquitoes to identify the salivary gland related proteins (Figure 3). We observed that among the 3,790 proteins identified in *Ae. aegypti* heads, 830 proteins were previously known to be expressed in the salivary glands. Moreover, we identified 77% of the proteins described by Wasinpiyamongkol et al. (2010) and 65% by Dhawan et al. (2017) in previous salivary gland proteome analysis. The 16 salivary proteins that were detected in all datasets probably are the most expressed and include inhibitors of platelet aggregation (salivary apyrase, AAEL006333-PA; apyrase precursor, AAEL006347PA), clotting inhibitors (SRPN23, AAEL002704-PB; SRPN26, AAEL003182-PA), aegyptins (AAEL010228-PA, AAEL010235-PA), angiopoietins (AAEL000749-PA, AAEL000726-PA), D7 family proteins (AAEL006417-PA, AAEL006424-PA), among others. We compared the 830 proteins previously known to be expressed in the salivary glands with the 196 differentially expressed proteins, to understand if these proteins could play a role during ZIKV and *Wolbachia* infections and/or co-infections. The modulation of saliva proteins is discussed below where the systemic changes in W, Z and WZ will be addressed.

**Figure 3.**
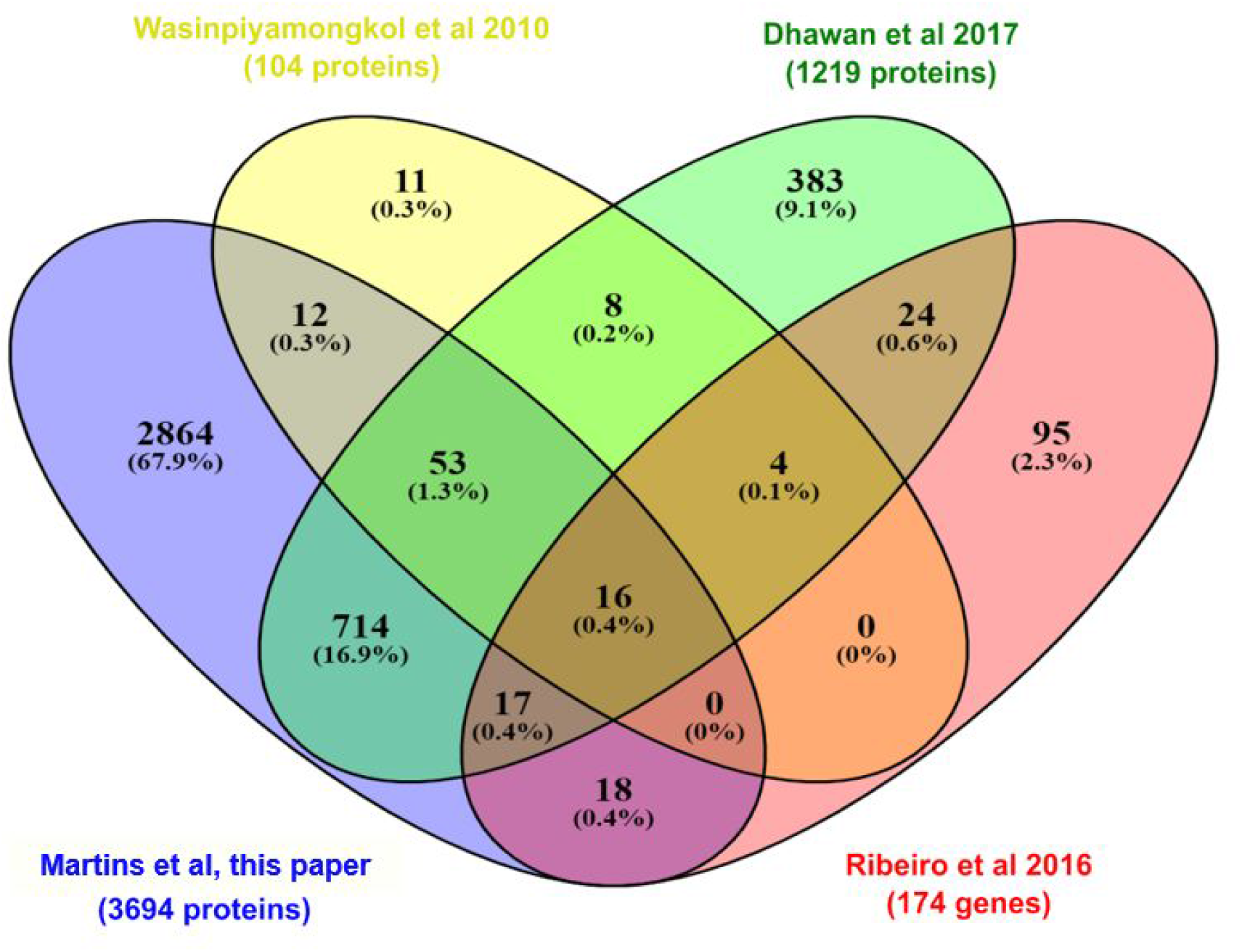
Venn diagram comparing the set of proteins (isoforms were not considered) identified in this work (purple) with two previous proteomes of the salivary glands of *Ae. aegypti* (yellow; Wasinpiyamongkol et al 2010 and green; Dhawan et al 2017) and one transcriptome (red; Ribeiro et al 2016) also carried out in salivary glands of *Ae. aegypti*, but in non-infected females.

As a class of neuronal signal molecules, neuropeptides are produced by the neurosecretory cells, that are mainly located in the brain, suboesophageal ganglion, among others (Li et al., 2020). In insects, neuropeptides and their receptors play important physiological processes such as development, reproduction, behavior and feeding (Holt et al., 2002). A previous study that investigated the presence of neuropeptidomics in *Ae. aegypti* proteome carried out in the central nervous system (CNS) and midgut samples identified 43 neuropeptides and hormones (Predel et al., 2010). Of these, we were able to identify 13 in our head proteomics analysis (Figure 4): Pyrokinin (PK) (AAEL012060), Short Neuropeptide F (sNPF) (AAEL019691), Neuropeptide-like precursor-1(NPLP) (AAEL012640), Calcitonin-like diuretic hormone (DH-31) (AAEL008070), SIFamide (AAEL009858), Crustacean cardioactive peptide (CCAP) (AAEL000630), Allatotropin (AAEL009541), CAPA (AAEL005444), Sulfakinin (SK) (AAEL006451), Ovary Ecdysteroidogenic Hormone (OEH; Neuroparsin A homolog) (AAEL004155), Insulin-like peptide 1 and 7 (AAEL000937; AAEL024251), and Adipokinetic hormone 1 (AKH) (AAEL011996). A few neuropeptides and hormones were modulated by co-infection and will be discussed afterwards.

**Figure 4.**
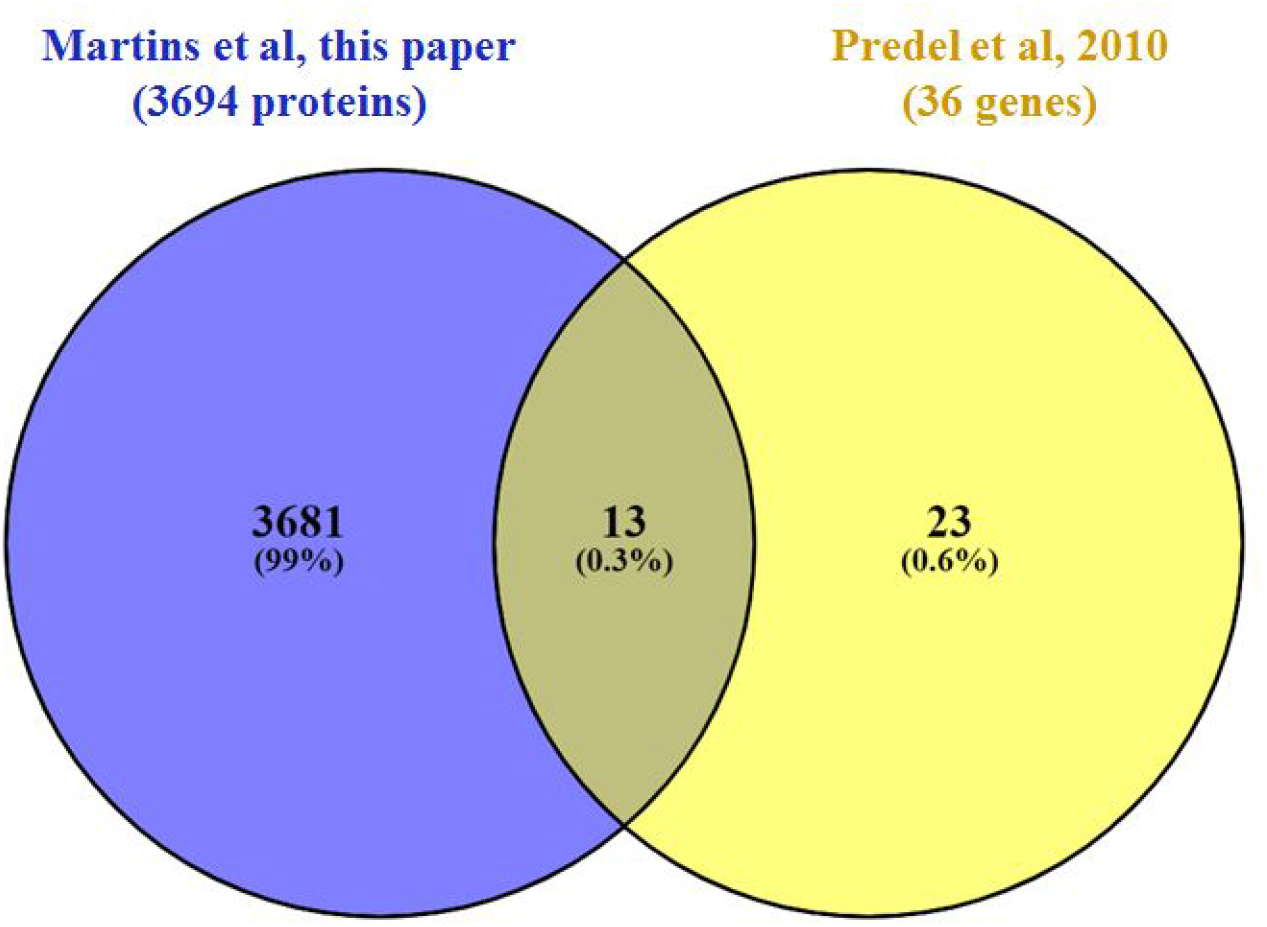
Venn diagram comparing the set of neuropeptides and hormones (isoforms were not considered) identified in this work (purple) with those identified in Predel (yellow; Predel et al 2010).

### 3.3 ZIKV infection modulates aerobic metabolism, lipids pathways, and immune system response

By comparing mosquitoes infected and non-infected with ZIKV (Figure 5; Supplementary Figure 4A), we observe four main enriched processes: nicotinamide nucleotide metabolism, lipid transport, humoral innate response and carbohydrate catabolism. Nicotinamide mononucleotide (NMN) is known for being an intermediator in NAD+ biosynthesis (Poddar et al., 2019; Hong et al., 2020), which participates actively in the aerobic metabolism process. This enrichment is mediated by the phosphoglycerate kinase (AAEL004988) and transaldolase (AAEL009389) upregulation, leading to carbohydrate catabolism elevation. The pentose phosphate pathway (PPP) was also up-regulated in our study as a response to transaldolase (AAEL009389) protein. This led us to relate the activation of PPP with the probation of nucleotide biosynthesis for virus replication (Guo et al., 2019). Both increased fluxes have been observed following Human Cytomegalovirus, Mayaro and Kaposi-sarcoma viruses in human cells (El-Bacha and Da Poian, 2013).

**Figure 5:**
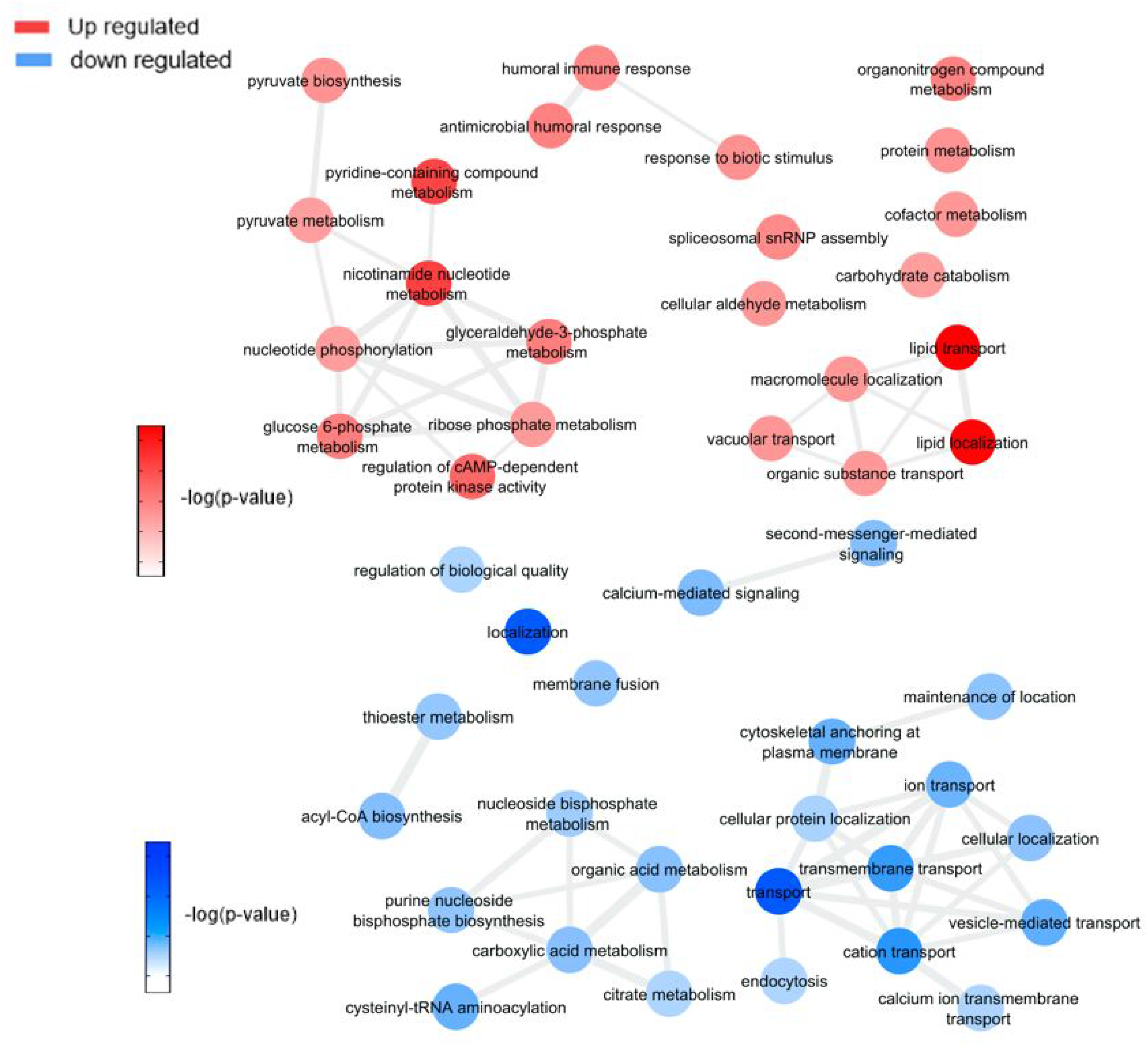
Overview of Significantly enriched GO [−log10 (P-value)] terms of ontological interaction network in biological processes of up-regulated (red) and down-regulated (blue) genes in ZIKV infected versus non-infected mosquitoes.

Viruses are capable of modulating mitochondrial bioenergetics aiming to synthesize macromolecular precursors for self-replication, assembly and egress (El-Bacha and Da Poian, 2013). Thus, the enhancement of the lipid transport process should drive a redirection of these energetic resources to the viral capsids formation (Kuzmina et al., 2020), an event that was also described in *Ae. aegypti* coinfected with DENV and *Wolbachia* (Koh et al., 2020). Koh et al. (2020) compared lipid abundances between non-infected and DENV-3 infected mosquitoes and found that most modulated lipids are glycerophospholipids, as described earlier (Perera et al., 2012), with a higher level of unsaturated fatty acids and triacylglycerols. Glycerophospholipids modulation can be related to a virus attempt to build new virus capsids, as these lipids are a compound of this structure (Martín-Acebes et al., 2016). In our results, three genes related to lipid localization and transport were up-regulated: apolipophorin-III (AAEL008789) and two vitellogenin-A1 precursors (AAEL006126, AAEL006138). Apolipophorins are protein components of lipoprotein particles that are essential for lipid transportation in the insect body, but apolipophorin-III can work as a pattern recognition receptor for insect immune system response (Wang et al., 2019), including in *Ae. aegypti* (Phillips and Clark, 2017). Lipid carrier protein lipophorin (Lp) and its lipophorin receptor (LpRfb) were significantly increased in *Ae. aegypti* after infections with Gram (+) bacteria and fungi, suggest that lipid metabolism is involved in the mosquito systemic immune responses to pathogens and parasites (Cheon et al., 2006). The apolipophorin-III upregulation can be rolled to lipid transport and also to the immune response as ZIKV is expected to regulate them and the protein participates in both processes.

Another group of up-regulated biological processes includes an innate immune response in host cells due to virus infection (antimicrobial humoral response, humoral immune response, and response to biotic stimulus). Insect immune system is generalist (https://doi.org/10.3389/fimmu.2017.00539), which may explain the identification of bacteria infection response pathways. Xi et al. (2008) found a significant role for the Toll pathway in DENV resistance regulation, and it was possible to identify differentially expressed antimicrobial peptide cecropin (AAEL029038) related to this pathway in our samples, confirming that Toll-pathway plays a major part in virus response (Xi et al., 2008; Pan et al., 2012; Kingsolver et al., 2013).

We observed down-regulation of tomosyn (AAEL006948), adenylate cyclase (AAEL009314), putative syntaxin-7 (AAEL009398), putative sodium/dicarboxylate cotransporter (AAEL009863), cytochrome c oxidase subunit II (AAEL018664), major facilitator superfamily domain-containing protein (AAEL006002), nuclear pore complex protein Nup188 (AAEL008557) and talin-1 (AAEL024235) related to cation transport processes. Also, three proteins were down-regulated: calcium-transporting ATPase sarcoplasmic/endoplasmic reticulum type (AAEL006582), Na+/K+ ATPase alpha subunit (AAEL012062), and calmodulin (AAEL012326-PA) related to cation transport processes as well. Similarly cytoskeleton, vesicle-mediated transport and ion transport-associated proteins down-regulation were described in CHIKV infection (Cui et al., 2020). Also, a recent study showed that Na+/K+ ATPase inhibition using two distinct drugs in mice helped to block ZIKV replication (Guo et al., 2020), suggesting that this down-regulation could be a mosquito cellular response to fight infection.

ATP-citrate synthase (AAEL004297), sphingosine phosphate lyase (AAEL003188) and probable cysteine-tRNA synthetase (AAEL022886), proteins associated with amino acids insertion in the citric acid cycle, were also down-regulated and may lead to a decrease in citric acid cycle and oxidative phosphorylation processes. This phenomenon was also observed in *Ae. aegypti* salivary glands infected with DENV-2 (Chisenhall et al., 2014; Shrinet et al., 2018), suggesting the occurrence of cellular biomolecule and energy production control by virus infection to avoid mitochondrial exhaustion.

### 3.4 *Wolbachia* endosymbiosis interfere in glycoconjugates production pathways, aerobic metabolism, immune response induction, and blood-feeding process

It is possible to notice in our data (Figure 6; Supplementary Figures 3B and 4B) an enrichment of GDP-mannose metabolism comparing mosquitoes non-infected with *Wolbachia*-mono infected. The enrichment was based on the upregulation of gdp mannose-4,6-dehydratase (AAEL014961), which converts GDP-mannose to GDP-4-dehydro-6-deoxy-D-mannose, the first of three steps for the conversion of GDP-mannose to GDP-fucose. This result suggests an induction caused by the presence of *Wolbachia* in insect cells to generate glycoconjugates, as GDP-mannose is a precursor for these molecules (Rexer et al. 2020; Stick and Williams, 2009), that may be used as elements to form new bacterial structures (García-del Portillo, 2020). An important observation was the enrichment of processes downregulated related to the citric acid cycle in mosquito cells infected with *Wolbachia*. As shown by Fernandes et al. (2014), *Wolbachia* modulated glycogen metabolism in *Ae. fluviatilis* embryos in favor of its aerobic metabolism (Fernandes et al., 2014).

**Figure 6:**
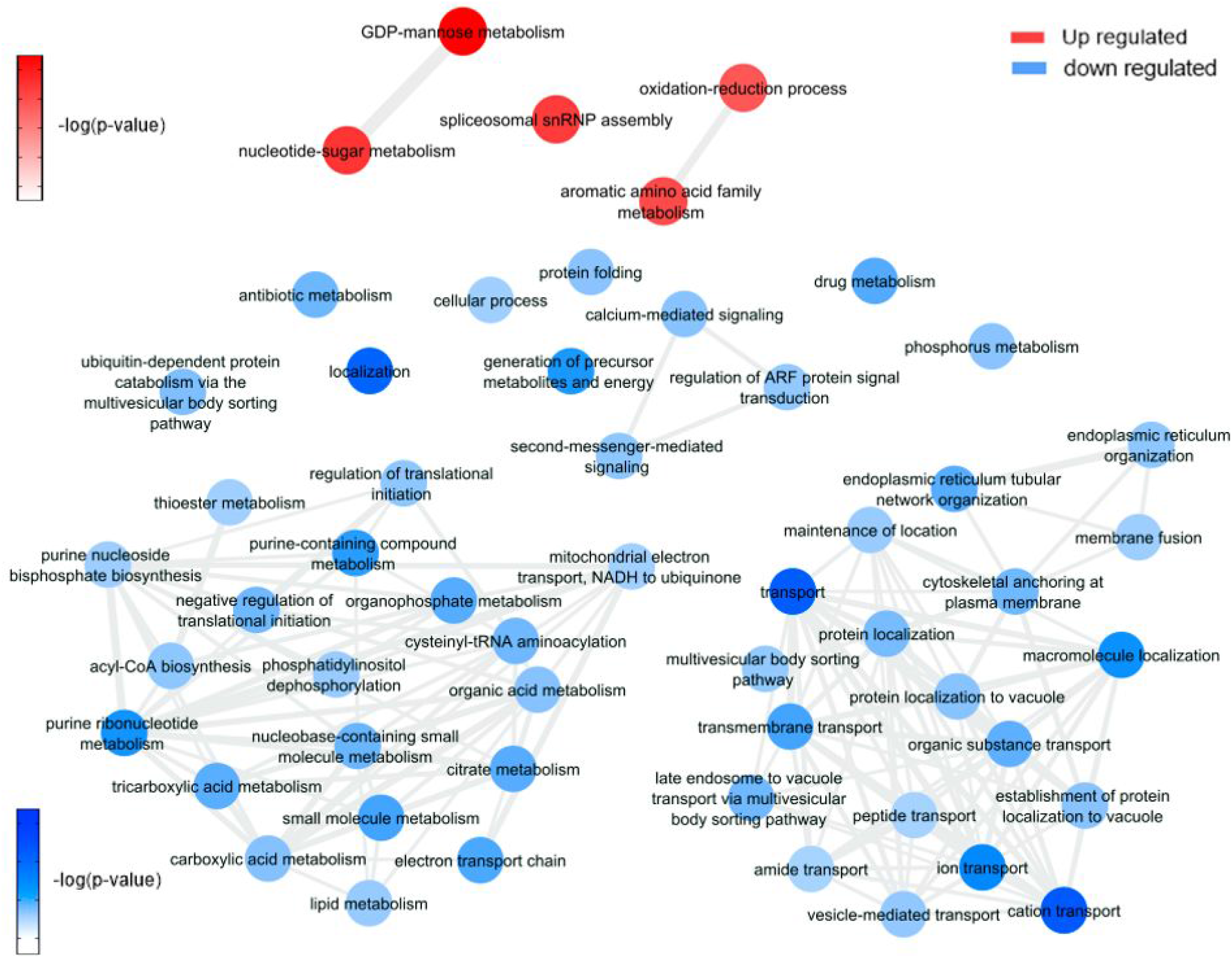
Overview of Significantly enriched GO [−log10 (P-value)] terms of ontological interaction network in biological processes of up-regulated (red) and down-regulated (blue) genes in *Wolbachia* infected versus non-infected mosquitoes.

Up-regulation of 4-hydroxyphenylpyruvate dioxygenase (AAEL014600) can be related to plastoquinol and vitamin E (tocopherol) synthesis. Both molecules are described in the literature as participants of antioxidant response to different stimuli in organisms (MaitiDutta et al., 2020; Nowicka et al., 2020), including oxidative stress (Kumar et al., 2020; Minter et al., 2020). In this way, we consider that genes correlated to these pathways were regulated during *Wolbachia* infection because this microorganism can induce the expression of antioxidant proteins (Brennan et al., 2008). Moreover, 4-hydroxyphenylpyruvate dioxygenase is related to L-tyrosine degradation I and L-phenylalanine degradation IV, which are essential traits for blood-feeding arthropods, as they consume an amount of blood proteins and need to deal with a high quantity of free amino acids (Sterkel et al., 2016).

The *Wolbachia* infection up-regulated 10 salivary gland proteins, such as endoplasmin (AAEL012827-PA), which is required for proper folding of Toll-like receptors in mammals (Fitzgerald and Kagan, 2020), and FKBP12 (AAEL010491-PB) that has a prolyl isomerase activity and bidins to immunosuppressive molecules (Caminati, and Procacci, 2020). 20 salivary proteins were down-regulated by *Wolbachia* infection. Interestingly, the same ion related proteíns down-regulated by ZIKV are also down-regulated by *Wolbachia*, moreover, proteins related with contraction such as paramyosin (AAEL010975-PA), tropomyosin (AAEL002761-PA), and Actin-1 (AAEL001928-PA) were also down-regulated. Also, a Serine protease inhibitor 25 (SRPN25) (AAEL007420-PB) was also identified in our analysis. Its orthologue is Alboserpin, the major salivary gland anticoagulant in *Ae. albopictus*, which prevents coagulation in an atypical reversible interaction with factor Xa (coagulation activation factor), important for blood intake by female mosquitoes (Stark and James, 1998; Calvo et al., 2011). SRPN25 was up-regulated in W, which may infer that blood takes longer to coagulate, increasing mosquito enzymes access to digest blood. The consequence of these salivary proteins modulation suggests an impairment in salivary gland injection into the host and a possible reduction in the amount of blood ingestion during *Wolbachia* presence in the mosquito (Turley et al., 2009).

### 3.5 Co-infection modulated pathways and processes show a reactive oxygen species accumulation leading to an increase in immune system response to virus infection

This analysis was based on a comparison of the ZW with Z (Figure 7; Supplementary Figure 4C) and W (Figure 8; Supplementary Figure 4D) mosquito heads samples.

**Figure 7:**
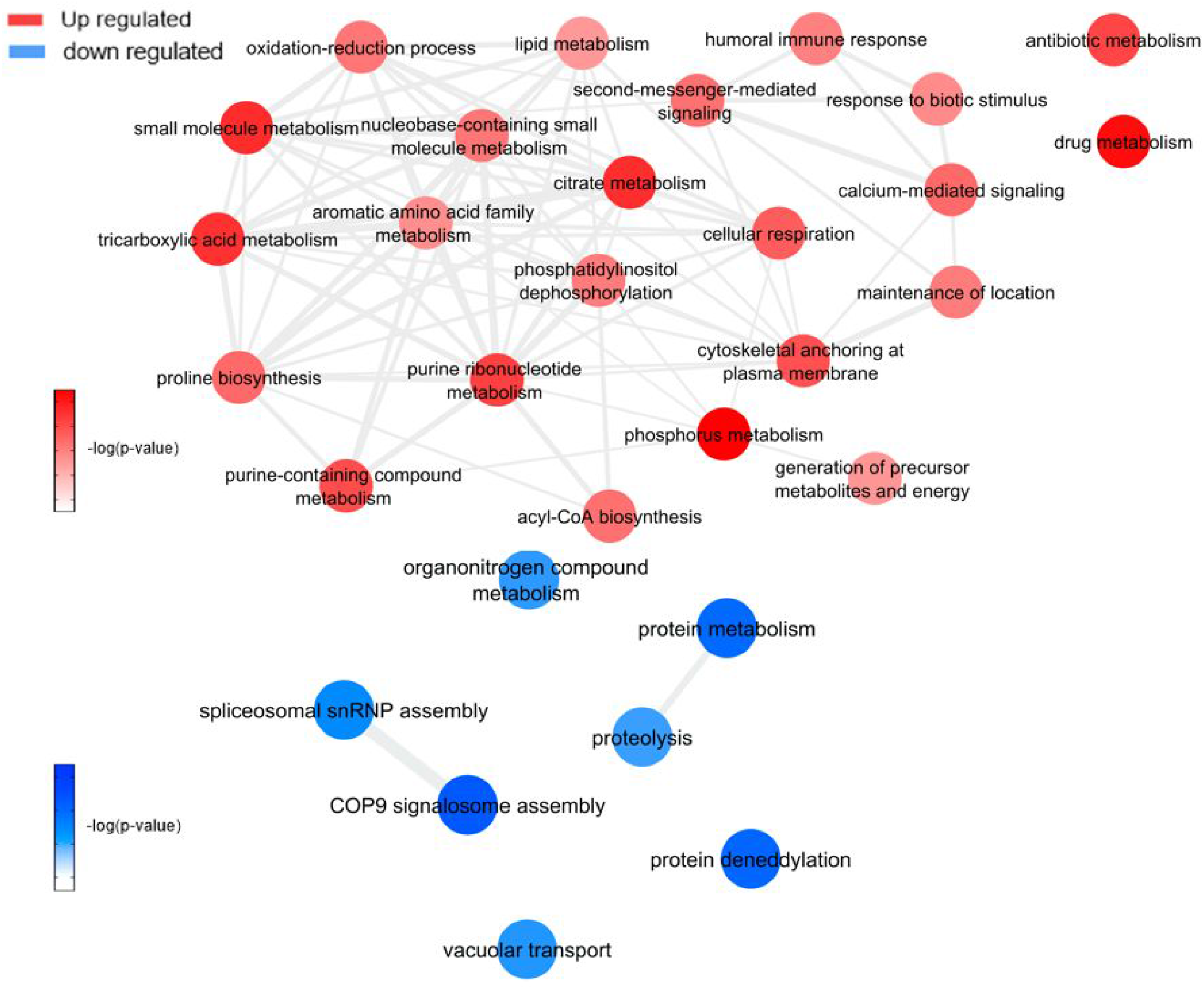
Overview of Significantly enriched GO [−log10 (P-value)] terms of ontological interaction network in biological processes of up-regulated (red) and down-regulated (blue) genes in coinfected mosquitoes versus ZIKV infected mosquitoes.

**Figure 8:**
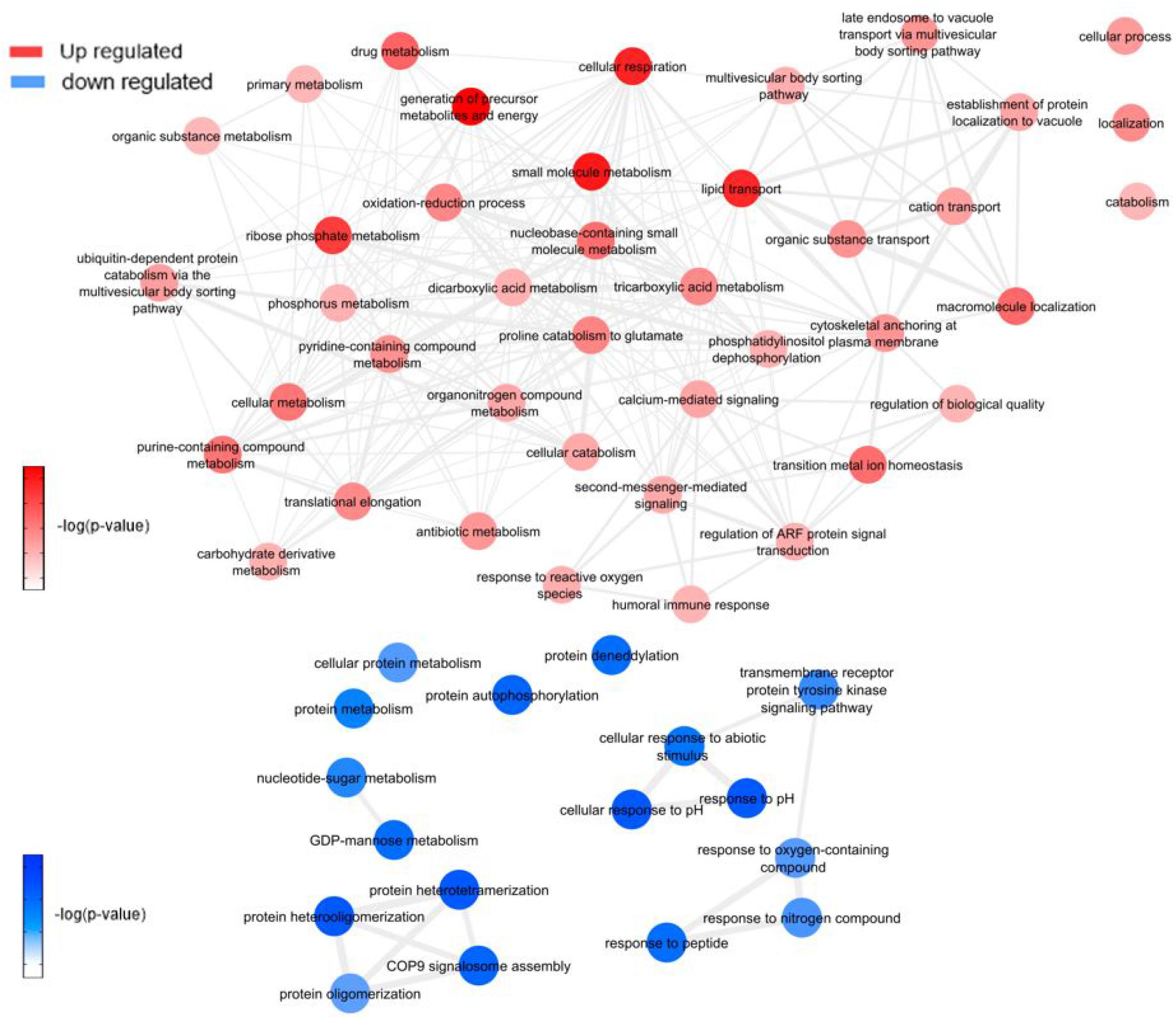
Overview of Significantly enriched GO [−log10 (P-value)] terms of ontological interaction network in biological processes of up-regulated (red) and down-regulated (blue) genes in coinfected mosquitoes versus *Wolbachia* infected mosquitoes.

ZIKV and *Wolbachia* use mosquitoes cell machinery respectively to synthesize lipids and glycoconjugates molecules for their structural use, as mentioned before. According to WZ results, the enriched lipid synthesis biological processes remain up-regulated despite the *Wolbachia* presence, similarly to ZIKV infection. Koh et al. (2020) confirmed such behavior during co-infection-induced intracellular events with DENV and *Wolbachia*, suggesting that they do not compete by lipids in host cell and virus-driven modulation dominates over that of *Wolbachia* (Koh et al., 2020), which may have occurred as well as in our experiment. In contrast, there is a down-regulation in GDP-mannose proteins genes synthesis, which may suggest that host cells no longer produce glycoconjugates for bacteria use.

Within the up-regulated genes in WZ compared to Z, ATP synthesis (ATPase subunit O; AAEL010823-PA), and cytoskeleton dynamics like microtubule binding and associated-proteins (AAEL009375-PK, AAEL004176-PB and AAEL009847-PB) suggest activation of the ubiquitin-proteasome system. Indeed, it was previously reported in *Drosophila* and mosquito cell lines transfected, that *Wolbachia* relies on host proteolysis through ubiquitination to acquire essential amino acid for the infection (Fallon and Witthuhn, 2009; White et al., 2017). Flavivirus also need a functioning ubiquitin-proteasome system to complete their life cycle (Choy et al., 2015a, 2015b), however, Ub3881 (AAEL003881), a ubiquitin-protein, is highly down-regulated in mosquitoes infected with Dengue virus (DENV) and its overexpression was able to control the virus infection (Troupin et al., 2016). In our analysis, we identified a paralog of the Ub3881 ubiquitin protein (AAEL003888), which was up-regulated in WZ vs. Z and down-regulated in Z, suggesting that *Wolbachia* have a role in AAEL003888 expression enhancement which might target the envelope proteins of ZIKV for degradation, therefore hindering the virus assembly.

Exploring neuropeptides and hormones, neuropeptide Sulfakinin (SK) (AAEL006451-PB) and Insulin-like peptide (ILP) 1 (AAEL000937-PA) were down-regulated comparing WZ with Z. SK are a family of neuropeptides, homologous to mammalian gastrin/cholecystokinin (CCK), responsible for food uptake control in insects (Zels et al., 2015) while ILPs are known for signaling carbohydrates intake in cells (Nässel and Broeck, 2016). A recent study correlated SK influence in ILPs concentration, where a rise in SK concentration promotes ILPs production (Słocińska et al., 2020). SK down-regulation, observed in our data, may decrease in mosquito feeding satisfaction. This fact allied to ILP down-regulation, leading to a decrease in glycolysis cell assimilation, can be understood as a virus-infection influence to increase mosquito blood uptake, elevating ZIKV transmission success.

There are several processes up-regulated after enrichment comparison between WZ and Z. Cellular respiration, aerobic respiration, small molecule metabolic process, tricarboxylic acid cycle, oxidative phosphorylation, ATP synthesis coupled electron transport, respiratory electron transport chain, generation of precursor metabolites and energy, tricarboxylic acid metabolic process, nicotinamide nucleotide metabolic process and cellular process were enriched up-regulated biological process by proteins genes ATP-citrate synthase (AAEL004297), glutamate semialdehyde dehydrogenase (AAEL006834), ATP synthase delta chain (AAEL010823), phosphatidylinositol-4,5-bisphosphate 4-phosphatase (AAEL011216), NADH-ubiquinone oxidoreductase (AAEL012552), mitochondrial aconitase (AAEL012897), 4-hydroxyphenylpyruvate dioxygenase (AAEL014600), titin (AAEL002565) and myosin (AAEL000596) up-regulation. The increase in aerobic respiration metabolism in *Wolbachia* infection proved to stimulate ROS production in insect cells as part of the host immune response, but it counterbalances with antioxidant pathways activation (Zug and Hammerstein, 2015). However, it was observed in WZ that the aerobic metabolism increases together with transition metal ions biological process and pathways, mediated by putative secreted ferritin G subunit precursor (AAEL007383) and transferrin (AAEL015458), which can also generate ROS (Liu et al., 2016). ROS likely activates the Toll-immune pathway to confront ZIKV, similar to DENV infection (Selivanov et al., 2008). Also, two eukaryotic elongation factor 1-alpha (eEF1A) proteins (AAEL017096; AAEL017301) were up-regulated comparing WZ and Z. eEF1A increase was described as a response to endoplasmic reticulum stress caused by ROS in Chinese hamster ovary (Borradaile et al., 2005), which may have occurred in the mosquito. Another observed data was SRPN25 down-regulated in WZ, which was up-regulated in W. SRPN25 down-regulation in WZ highlights its function related to the insect immune system. Serpins can block the Toll signaling cascade and β-1,3-glucan-mediated melanin biosynthesis (Jiang et al., 2009), so its downregulation can infer that the insect immune system is activated.

This led us to understand insects defense mechanisms, induced by the presence of endosymbiotic bacteria, that may have been acting against virus infection: melanization activation mediated by SRPN25 down-regulation and Toll-pathway with the influence of SPRN25 down-regulation and the exacerbated production of reactive oxygen species (ROS) by aerobic metabolism increase. ROS occurs inside the mitochondria, after the oxidative phosphorylation stage, which has oxygen as the final electron acceptor, forming metabolic water. When oxygen is prematurely and incompletely reduced, it gives rise to superoxide anion, classified as ROS, together with hydrogen peroxide, hypochlorous acid, hydroxyl radical, singlet oxygen, and ozone (Lambert and Brand, 2009). Pan et al 2011 described the relation between *Wolbachia* presence in *A. aegypti* and Toll-pathway activation by ROS to control DENV (Pan et al., 2012). This study not only confirms oxidative stress caused by Wolbachia, but also antioxidant genes expression as described earlier in this paper. The role of symbiotic microorganisms in arbovirus infection of arthropods vectors is widely discussed, highlighting the importance of a ROS-mediated stimulation of the Toll-pathway and AMPs expression (Yin et al., 2020). As SPRN25 is down-regulated, serpins activity is up-regulated, which lead to serpins key response in the defense mechanism of insects, activating especially the Toll pathway, as well as ROS, and prophenoloxidase (PPO) cascade (Meekins et al., 2017). As for PPO cascade, it mediates melanization immune response in insects, including mosquito vectors (Christensen et al., 2005), It was already investigated that melanization suppression by SRPN activation can lead to a increase in viral infection (Toufeeq et al., 2019), calling attention to SRPN importance in mosquito immune response.

## 4 Conclusions

Proteomics analysis of female’s head of *Ae. aegypti* during mono-infection with ZIKV or *Wolbachia* highlighted how those microorganisms use cell host for their benefits. Our data support that ZIKV induces lipid synthesis and transport allied with glycolysis pathways while *Wolbachia* increases glycoconjugates production. During co-infection, *Wolbachia* helps *Ae. aegypti* to prevent virus infection by stimulating ROS production leading to Toll-pathway humoral immune response, together with antioxidant production to control cell homeostasis. This mechanism seems to be efficient since we have shown that peptides coming from the ZIKV polyprotein are reduced in female’s heads of *Ae aegypti* in the presence of *Wolbachia*. Our study has provided important insights into the differential manner in which *Ae. aegypti* proteome is systemically regulated during mono- and co-infections of ZIKV and *Wolbachia*. Ultimately, this work represents a rich resource for the insect research community to help elucidate the mechanisms by which *Wolbachia* orchestrate resistance to ZIKV infection in *Ae. aegypti*

## Availability of raw files

The mass spectrometry proteomics data have been deposited to the ProteomeXchange Consortium via the PRIDE partner repository with the dataset identifier PXD022665. **Username:** **Password:** wGLeEMiW

## Authors Contribution

MM, LFCR, JRM, AT and SSC performed experiments and data analysis. MM and SSC produced paper figures. MM, LFCR, JRM, AT, SSC, GBD, DMPO, RDM, FCSN, RM and MJ wrote the manuscript. RM and MJ idealized and coordinated the study. All authors approved the manuscript.

## Acknowledgements

The authors thank Fabio Mendonça Gomes from Instituto de Biofísica Carlos Chagas Filho - UFRJ and Livia Goto-Silva from D’Or Institute for Research and Education (IDOR) for their critical reading of the manuscript.

