## Supplementary material for "Comprehensive quantitative proteome analysis of *Aedes aegypti* identifies proteins and pathways involved in *Wolbachia pipientis* and Zika virus interference phenomenon": Suplemental Figures 1, 2, 3 and 4.

Supplementary figure 1 - A: MS / MS spectra of the LITANPVITESTENS peptide of the ZIKV polyprotein B: MS / MS spectra of the SHTLWTDGIEESDLIIP peptide of the ZIKV polyprotein C: MS / MS spectra of the LITANPVITESTENS peptide of the ZIKV polyprotein

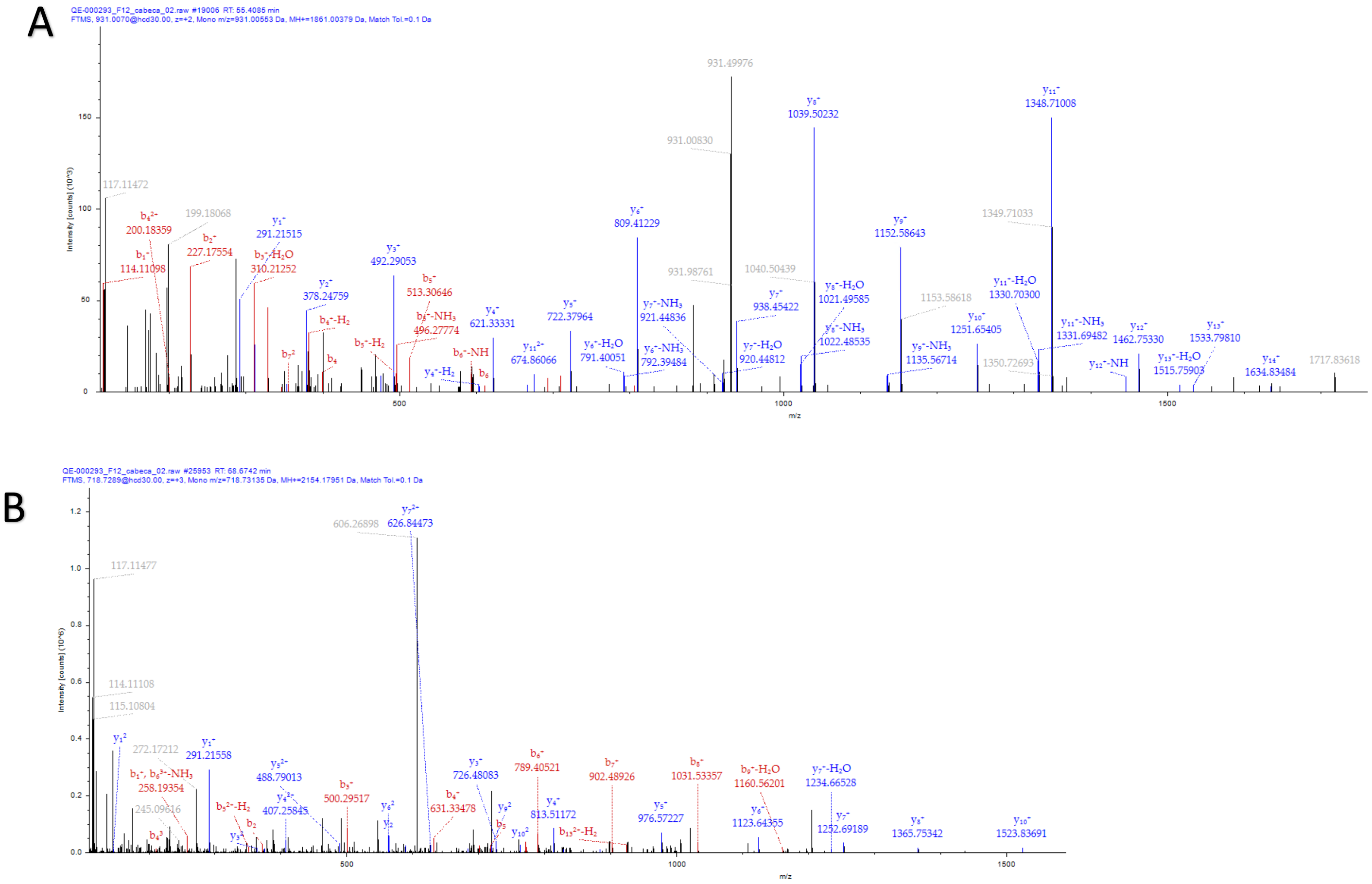

C

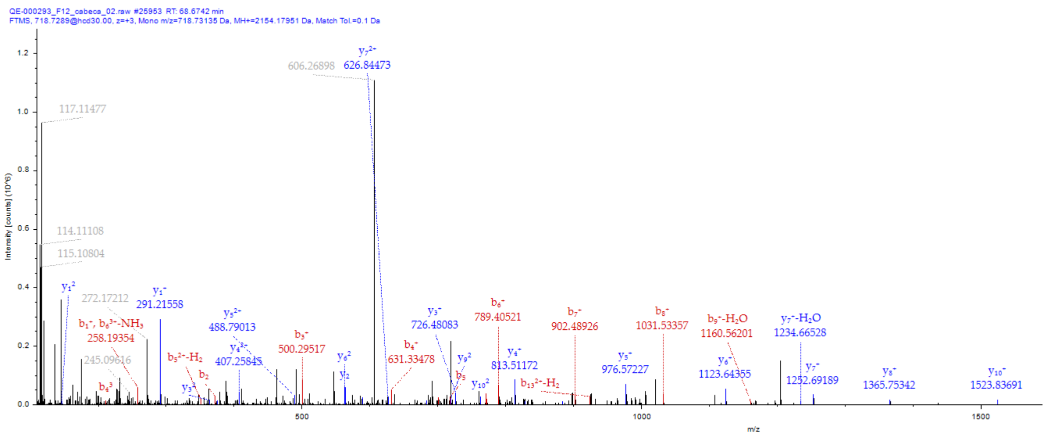


Supplementary figure 2 – Peptide wolbachia abundance identified.

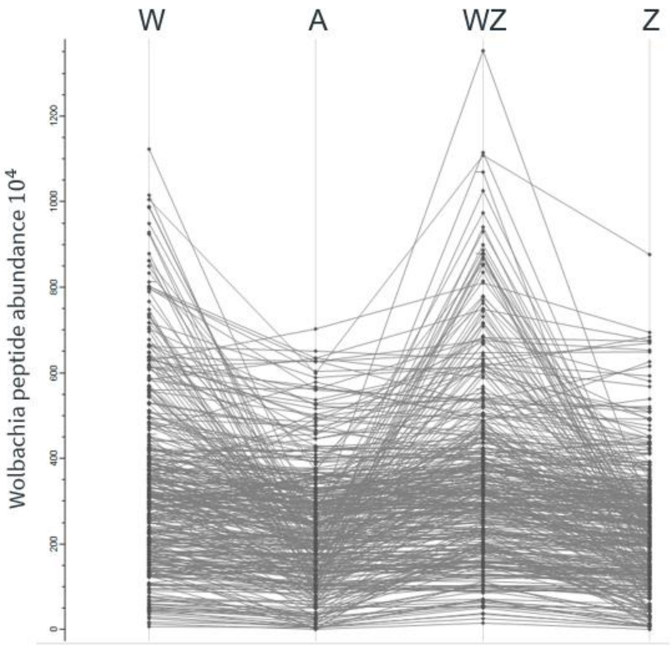

Supplementary figure 3 - A: Charts showing before and after normalization of proteins. B: histogram of each sample showing normal distributions of the intensities C: Pearson correlation coefficients between technical replicates of each sample.

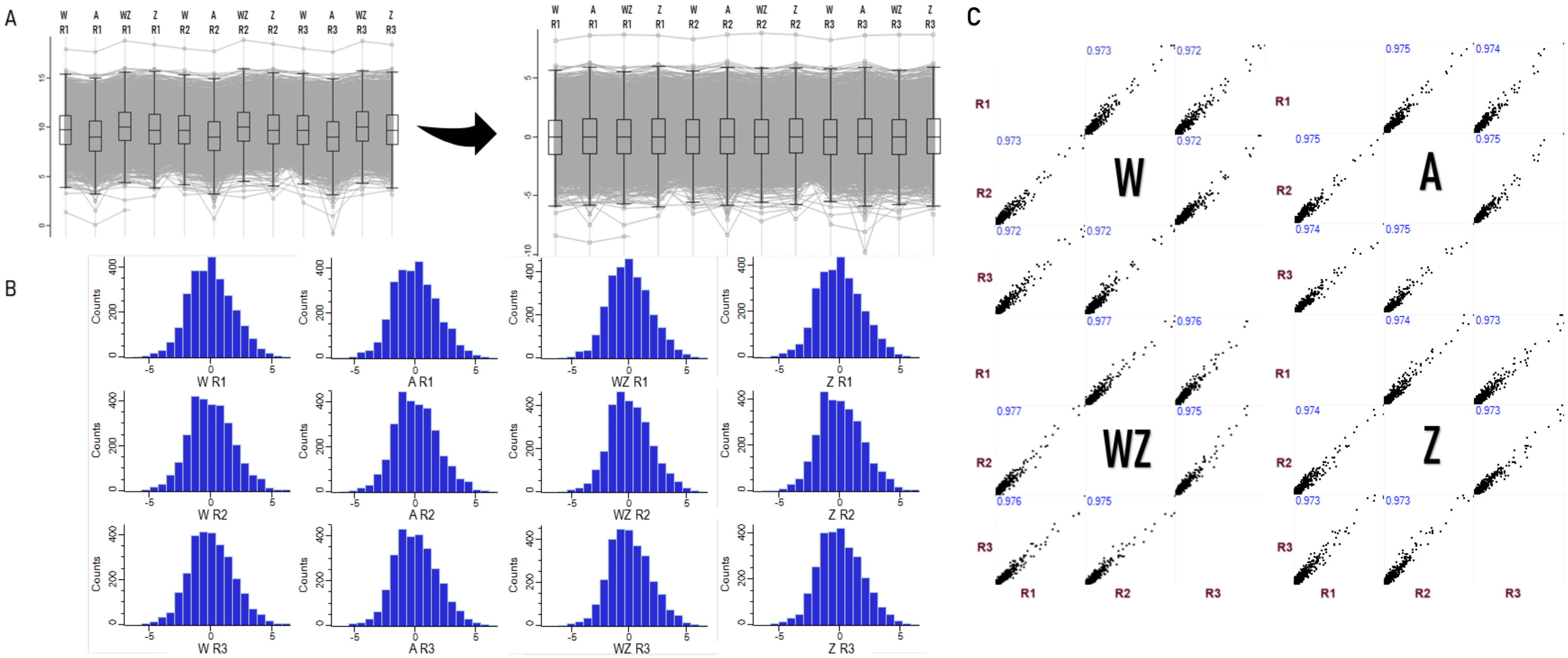

Supplementary figure 4 - Bar charts of biological processes (gene ontology terms) enriched analysis in A: ZIKV infected versus non-infected; B: *Wolbachia* infected versus non-infected mosquitoes. C: coinfected mosquitoes versus *ZIKV* infected mosquitoes D: coinfected mosquitoes versus *Wolbachia* infected mosquitoes.

A

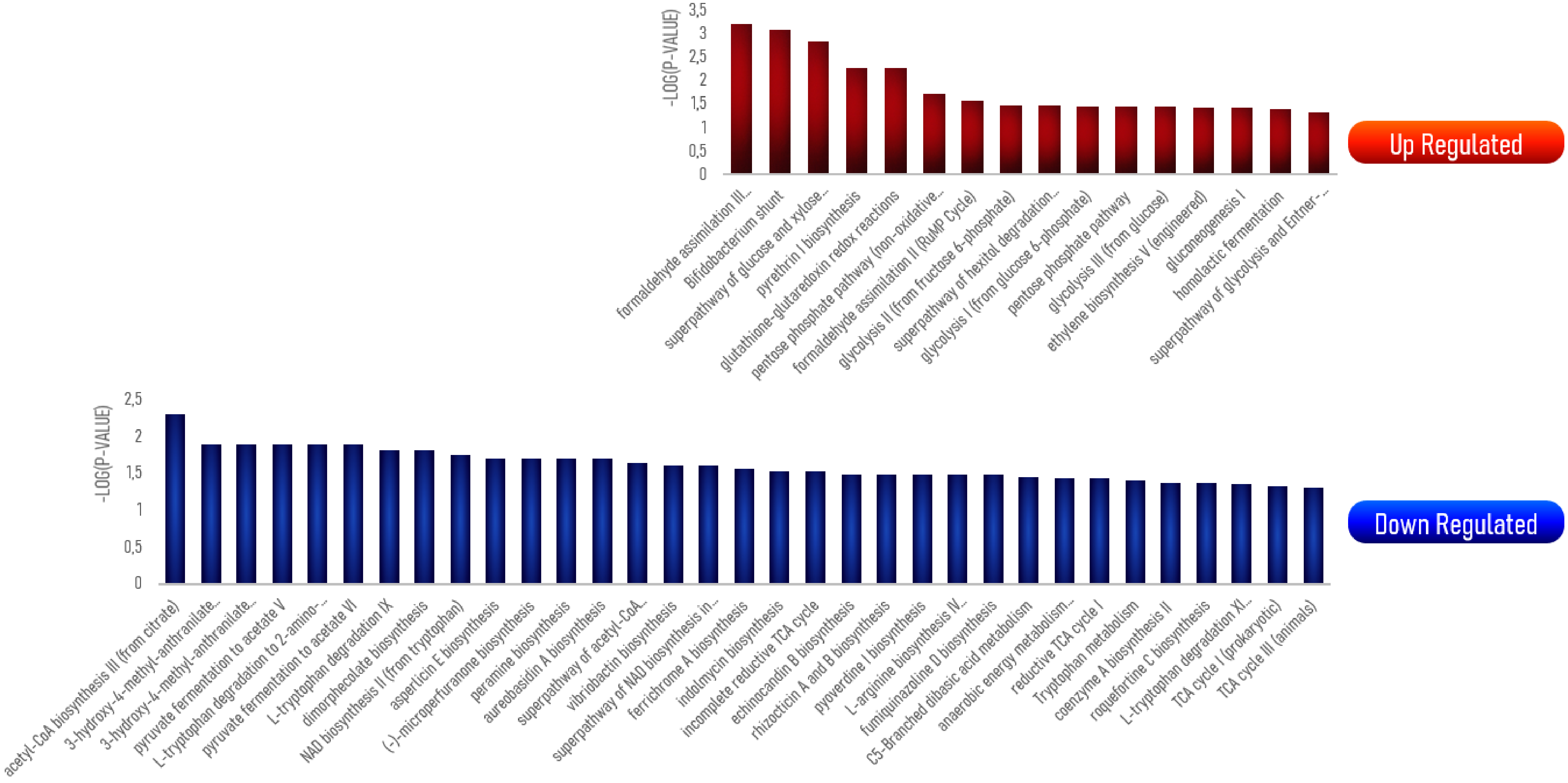

-LOG(P-VALUE)
3,5
3
2,5
2
1,5
1
0,5
0
formaldehyde assimilation III...
Bifidobacterium shunt
superpathway of glucose and xylose...
pyrethrin I biosynthesis
glutathione-glutaredoxin redox reactions
pentose phosphate pathway (non-oxidative...
formaldehyde assimilation II (RuMP Cycle)
glycolysis II (from fructose 6-phosphate)
superpathway of hexitol degradation...
glycolysis I (from glucose 6-phosphate)
pentose phosphate pathway
glycolysis III (from glucose)
ethylene biosynthesis V (engineered)
gluconeogenesis I
homolactic fermentation
superpathway of glycolysis and Entner-...
Up Regulated
-LOG(P-VALUE)
2,5
2
1,5
1
0,5
0
acetyl-CoA biosynthesis III (from citrate)
3-hydroxy-4-methyl-anthranilate...
3-hydroxy-4-methyl-anthranilate...
pyruvate fermentation to acetate V
L-tryptophan degradation to 2-amino-...
pyruvate fermentation to acetate VI
L-tryptophan degradation IX
dimorphecolate biosynthesis
NAD biosynthesis II (from tryptophan)
aspterricin E biosynthesis
(-)-microperfuranone biosynthesis
peramine biosynthesis
aureobasidin A biosynthesis
superpathway of acetyl-CoA...
vibriobactin biosynthesis
superpathway of NAD biosynthesis in...
ferrichrome A biosynthesis
indolmycin biosynthesis
incomplete reductive TCA cycle
echinocandin B biosynthesis
rhizocticin A and B biosynthesis
pyoverdine I biosynthesis
L-arginine biosynthesis IV...
fumiquinazoline D biosynthesis
C5-Branched dibasic acid metabolism
anaerobic energy metabolism...
reductive TCA cycle I
Tryptophan metabolism
coenzyme A biosynthesis II
roquefortine C biosynthesis
L-tryptophan degradation XI...
TCA cycle I (prokaryotic)
TCA cycle III (animals)
Down Regulated

B

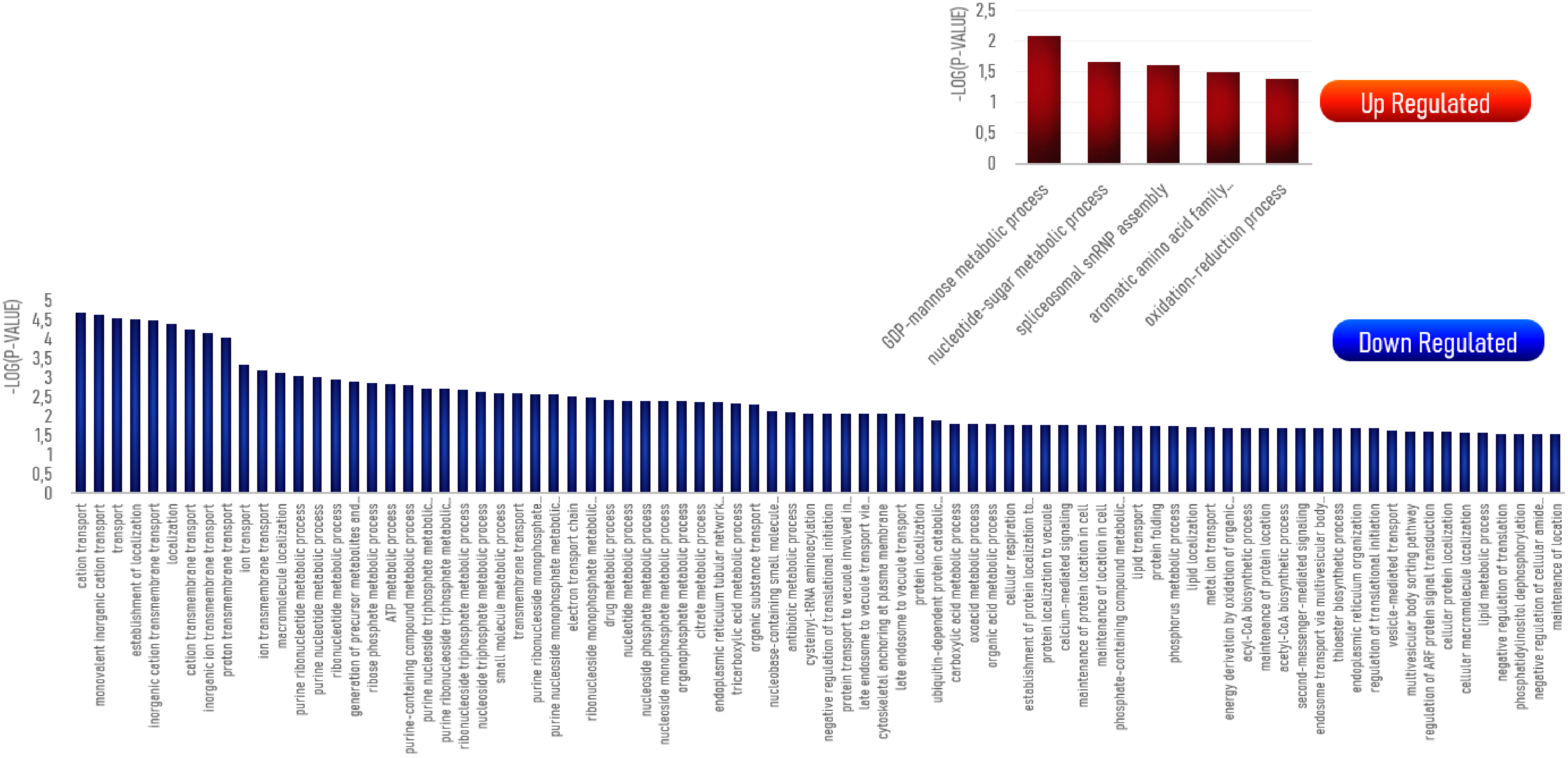
Up Regulated
-LOG(P-VALUE)
2,5
2
1,5
1
0,5
0
GDP-mannose metabolic process
nucleotide-sugar metabolic process
spliceosomal snRNP assembly
aromatic amino acid family...
oxidation-reduction process
Down Regulated
-LOG(P-VALUE)
5
4,5
4
3,5
3
2,5
2
1,5
1
0,5
0
cation transport
monovalent inorganic cation transport
transport
establishment of localization
inorganic cation transmembrane transport
localization
cation transmembrane transport
inorganic ion transmembrane transport
proton transmembrane transport
ion transport
ion transmembrane transport
macromolecule localization
purine ribonucleotide metabolic process
purine nucleotide metabolic process
ribonucleotide metabolic process
generation of precursor metabolites and...
ribose phosphate metabolic process
ATP metabolic process
purine-containing compound metabolic process
purine nucleoside triphosphate metabolic...
purine ribonucleoside triphosphate metabolic...
ribonucleoside triphosphate metabolic process
nucleoside triphosphate metabolic process
small molecule metabolic process
transmembrane transport
purine ribonucleoside monophosphate...
purine nucleoside monophosphate metabolic...
electron transport chain
ribonucleoside monophosphate metabolic...
drug metabolic process
nucleotide metabolic process
nucleoside phosphate metabolic process
nucleoside monophosphate metabolic process
organophosphate metabolic process
citrate metabolic process
endoplasmic reticulum tubular network...
tricarboxylic acid metabolic process
organic substance transport
nucleobase-containing small molecule...
antibiotic metabolic process
cysteinyl-tRNA aminoacylation
negative regulation of translational initiation
protein transport to vacuole involved in...
late endosome to vacuole transport via...
cytoskeletal anchoring at plasma membrane
late endosome to vacuole transport
protein localization
ubiquitin-dependent protein catabolic...
carboxylic acid metabolic process
oxoacid metabolic process
organic acid metabolic process
cellular respiration
establishment of protein localization to...
protein localization to vacuole
calcium-mediated signaling
maintenance of protein location in cell
maintenance of location in cell
phosphate-containing compound metabolic...
lipid transport
protein folding
phosphorus metabolic process
lipid localization
metal ion transport
energy derivation by oxidation of organic...
acyl-CoA biosynthetic process
maintenance of protein location
acetyl-CoA biosynthetic process
second-messenger-mediated signaling
endosome transport via multivesicular body...
thioester biosynthetic process
endoplasmic reticulum organization
regulation of translational initiation
vesicle-mediated transport
multivesicular body sorting pathway
regulation of ARF protein signal transduction
cellular protein localization
cellular macromolecule localization
lipid metabolic process
negative regulation of translation
phosphatidylinositol dephosphorylation
negative regulation of cellular amide...
maintenance of location

C

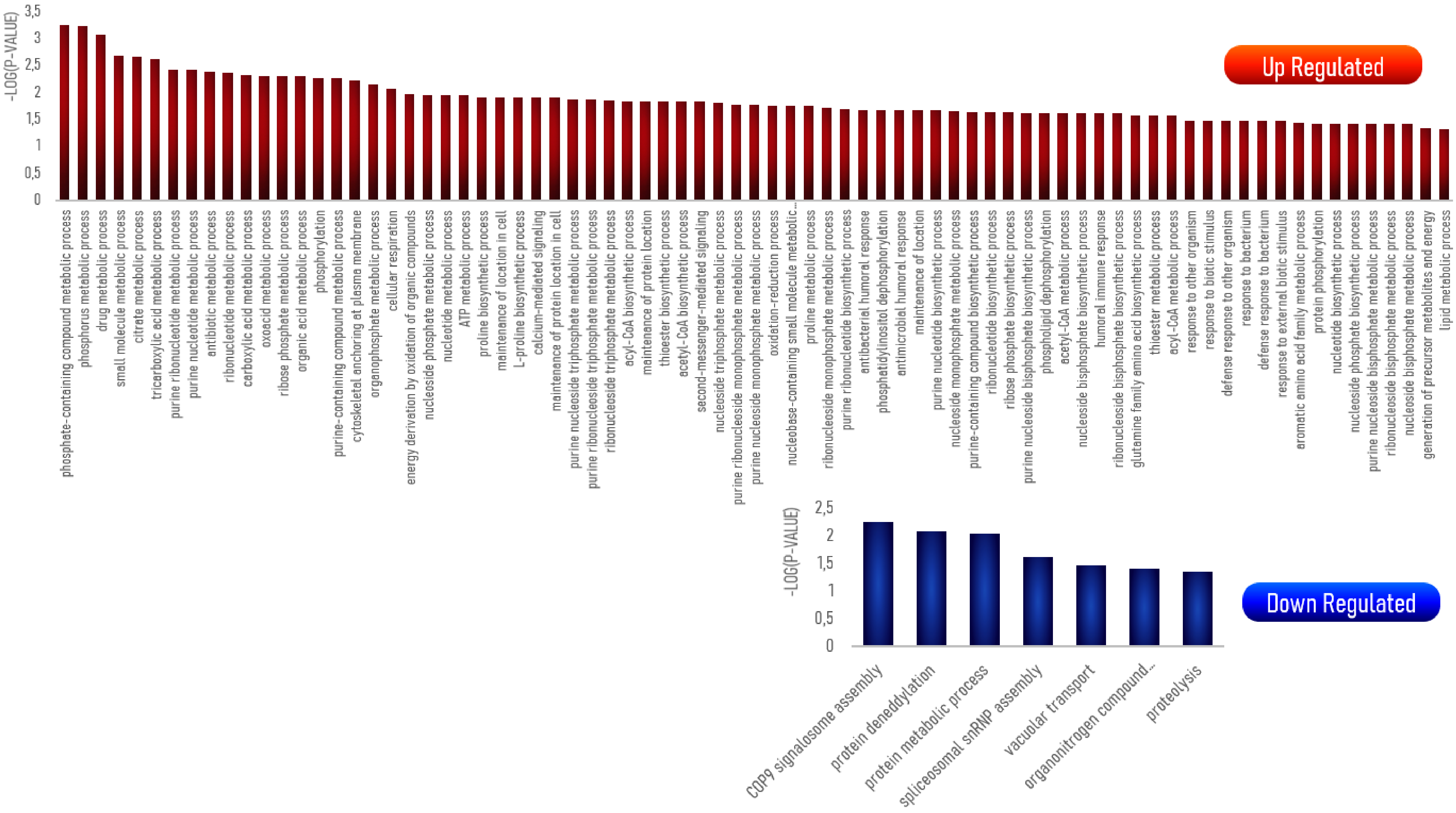
-LOG(P-VALUE)
3,5
3
2,5
2
1,5
1
0,5
0
Up Regulated
phosphate-containing compound metabolic process
phosphorus metabolic process
drug metabolic process
small molecule metabolic process
citrate metabolic process
tricarboxylic acid metabolic process
purine ribonucleotide metabolic process
purine nucleotide metabolic process
antibiotic metabolic process
ribonucleotide metabolic process
carboxylic acid metabolic process
oxoacid metabolic process
ribose phosphate metabolic process
organic acid metabolic process
phosphorylation
purine-containing compound metabolic process
cytoskeletal anchoring at plasma membrane
organophosphate metabolic process
cellular respiration
energy derivation by oxidation of organic compounds
nucleoside phosphate metabolic process
nucleotide metabolic process
ATP metabolic process
proline biosynthetic process
maintenance of location in cell
L-proline biosynthetic process
calcium-mediated signaling
maintenance of protein location in cell
purine nucleoside triphosphate metabolic process
purine ribonucleoside triphosphate metabolic process
ribonucleoside triphosphate metabolic process
acyl-CoA biosynthetic process
maintenance of protein location
thioester biosynthetic process
acetyl-CoA biosynthetic process
second-messenger-mediated signaling
nucleoside triphosphate metabolic process
purine ribonucleoside monophosphate metabolic process
purine nucleoside monophosphate metabolic process
oxidation-reduction process
nucleobase-containing small molecule metabolic...
proline metabolic process
ribonucleoside monophosphate metabolic process
purine ribonucleotide biosynthetic process
antibacterial humoral response
phosphatidylinositol dephosphorylation
antimicrobial humoral response
maintenance of location
purine nucleotide biosynthetic process
nucleoside monophosphate metabolic process
purine-containing compound biosynthetic process
ribonucleotide biosynthetic process
ribose phosphate biosynthetic process
purine nucleoside bisphosphate biosynthetic process
phospholipid dephosphorylation
acetyl-CoA metabolic process
nucleoside bisphosphate biosynthetic process
humoral immune response
ribonucleoside bisphosphate biosynthetic process
glutamine family amino acid biosynthetic process
thioester metabolic process
acyl-CoA metabolic process
response to other organism
response to biotic stimulus
defense response to other organism
response to bacterium
defense response to bacterium
response to external biotic stimulus
aromatic amino acid family metabolic process
protein phosphorylation
nucleotide biosynthetic process
nucleoside phosphate biosynthetic process
purine nucleoside bisphosphate metabolic process
ribonucleoside bisphosphate metabolic process
nucleoside bisphosphate metabolic process
generation of precursor metabolites and energy
lipid metabolic process
-LOG(P-VALUE)
2,5
2
1,5
1
0,5
0
Down Regulated
COP9 signalosome assembly
protein deneddylation
protein metabolic process
spliceosomal snRNP assembly
vacuolar transport
organonitrogen compound...
proteolysis

D

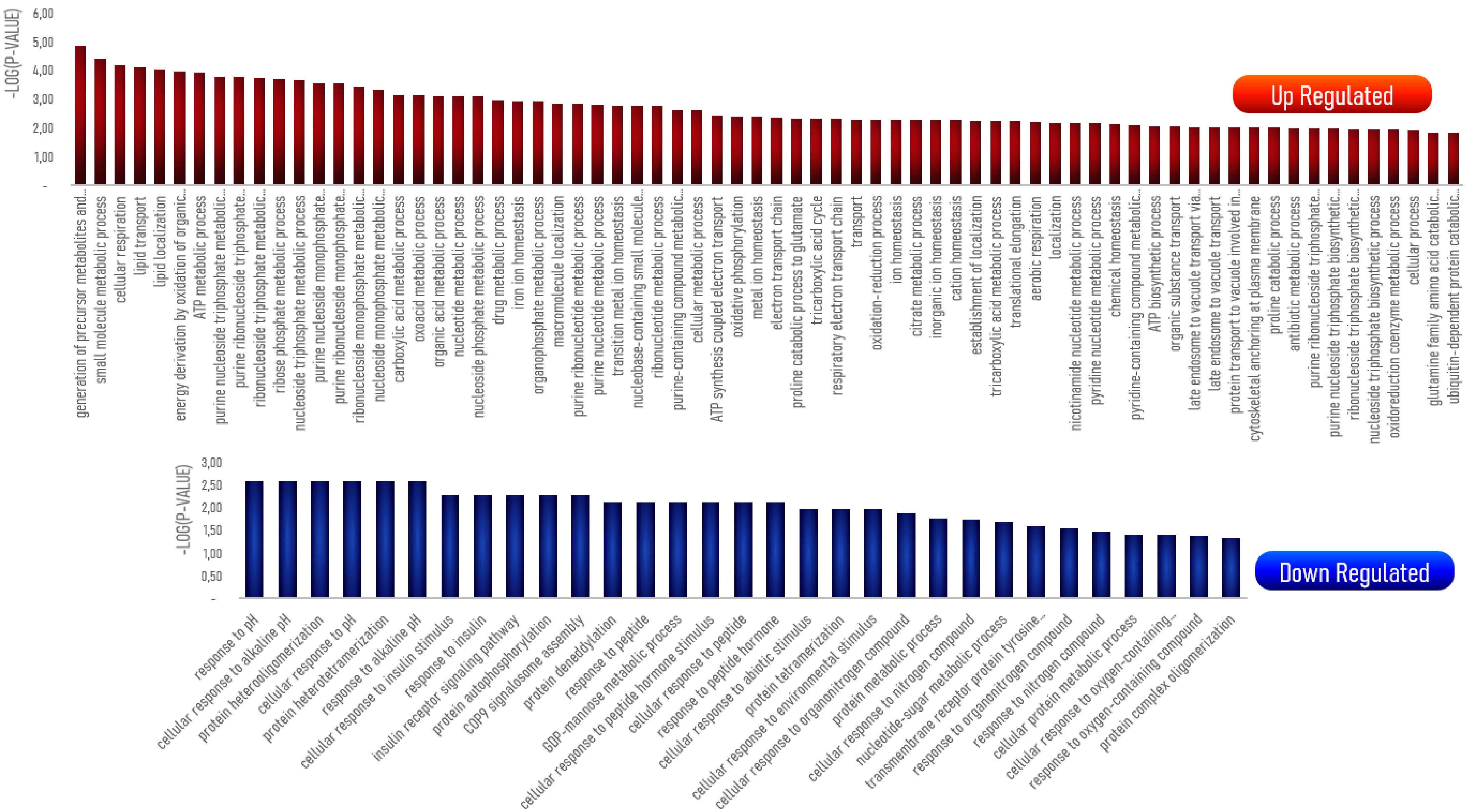

-LOG(P-VALUE)
6,00
5,00
4,00
3,00
2,00
1,00
-
Up Regulated
generation of precursor metabolites and...
small molecule metabolic process
cellular respiration
lipid transport
lipid localization
energy derivation by oxidation of organic...
ATP metabolic process
purine nucleoside triphosphate metabolic...
purine ribonucleoside triphosphate...
ribonucleoside triphosphate metabolic...
ribose phosphate metabolic process
nucleoside triphosphate metabolic process
purine nucleoside monophosphate...
purine ribonucleoside monophosphate...
ribonucleoside monophosphate metabolic...
nucleoside monophosphate metabolic...
carboxylic acid metabolic process
oxoacid metabolic process
organic acid metabolic process
nucleotide metabolic process
nucleoside phosphate metabolic process
drug metabolic process
iron ion homeostasis
organophosphate metabolic process
macromolecule localization
purine ribonucleotide metabolic process
purine nucleotide metabolic process
transition metal ion homeostasis
nucleobase-containing small molecule...
ribonucleotide metabolic process
purine-containing compound metabolic...
cellular metabolic process
ATP synthesis coupled electron transport
oxidative phosphorylation
metal ion homeostasis
electron transport chain
proline catabolic process to glutamate
tricarboxylic acid cycle
respiratory electron transport chain
transport
oxidation-reduction process
ion homeostasis
citrate metabolic process
inorganic ion homeostasis
cation homeostasis
establishment of localization
tricarboxylic acid metabolic process
translational elongation
aerobic respiration
localization
nicotinamide nucleotide metabolic process
pyridine nucleotide metabolic process
chemical homeostasis
pyridine-containing compound metabolic...
ATP biosynthetic process
organic substance transport
late endosome to vacuole transport via...
late endosome to vacuole transport
protein transport to vacuole involved in...
cytoskeletal anchoring at plasma membrane
proline catabolic process
antibiotic metabolic process
purine ribonucleoside triphosphate...
purine nucleoside triphosphate biosynthetic...
ribonucleoside triphosphate biosynthetic...
nucleoside triphosphate biosynthetic process
oxidoreduction coenzyme metabolic process
cellular process
glutamine family amino acid catabolic...
ubiquitin-dependent protein catabolic...
-LOG(P-VALUE)
3,00
2,50
2,00
1,50
1,00
0,50
-
Down Regulated
response to pH
cellular response to alkaline pH
protein heterooligomerization
cellular response to pH
protein heterotetramerization
response to alkaline pH
cellular response to insulin stimulus
response to insulin
insulin receptor signaling pathway
protein autophosphorylation
COP9 signalosome assembly
protein deneddylation
response to peptide
GDP-mannose metabolic process
cellular response to peptide hormone stimulus
cellular response to peptide
response to peptide hormone
cellular response to abiotic stimulus
protein tetramerization
cellular response to environmental stimulus
cellular response to organonitrogen compound
protein metabolic process
cellular response to nitrogen compound
nucleotide-sugar metabolic process
transmembrane receptor protein tyrosine...
response to organonitrogen compound
response to nitrogen compound
cellular protein metabolic process
cellular response to oxygen-containing...
response to oxygen-containing compound
protein complex oligomerization
